# Microscopic Elementary Current Properties Survive Across Scales Shaping Spectral And Non-Linear Features Of Field Potentials

**DOI:** 10.64898/2026.11.25.769229

**Authors:** Ricardo Muñoz-Arnaiz, Ana Ramírez-Frías, Faezeh Ashouri, Julia Makarova, Oscar Herreras

## Abstract

The temporal fluctuations of field potentials (FPs) carry information about upstream neuronal dynamics, but their extracellular expression is shaped by the anatomo-functional properties of the receiving population. Inferring upstream dynamics from FPs requires understanding how incoming activity is transformed by the receiving population. We here explored whether elementary synaptic current waveform shape contributes to mesoscopic fluctuations or is diluted during micro-to-mesoscopic integration. Using spatially stable hippocampal FP generators as templates for reconstruction provided pathway-specific temporal fluctuations while avoiding the ambiguities introduced by source geometry and mixing in raw FPs. Long epochs encompassing diverse temporal patterns helped identify general effects of elementary current properties. Synaptic-like unitary extracellular contributions (UECs) with prescribed waveforms were introduced, keeping the temporal sequence of microscopic events identical to isolate their impact on mesoscopic properties. UEC waveform shape influenced mean mesoscopic DC-like amplitude and both readily observable FP properties (amplitude, waveform and spectral composition) and less directly observable, predominantly non-spectral features captured by nonlinear, fractal and entropy-based measures. These effects were generator-specific, yet largely consistent across animals and independent surrogate event trains extracted from the original patterns. Thus, population integration does not erase the influence of elementary extracellular current properties, but transforms how they are expressed in mesoscopic FP features. These findings suggest that, in addition to upstream output patterns, postsynaptic contributions also shape both readily observable and less directly observable FP features, and that different pathways may contribute differently according to the properties of their elementary postsynaptic currents.

## INTRODUCTION

Despite the considerable attention devoted to recognizable and oscillatory patterns (Buzsáki et al., 2012), field-potential (FP) activity is predominantly irregular (Eckhorn, 1994; Bullock et al., 2003; Thivierge and Cisek, 2008; Dehghani et al., 2010; He et al., 2010). Its temporal and spectral properties reflect compound upstream dynamics, expressed through postsynaptic currents in receiving populations and variably shaped by the cellular and spatial organization of the receiving tissue (Martín-Vázquez et al., 2016). Indeed, only a subset of neuronal populations has the geometrical organization required for incoming synaptic activity to generate a measurable mesoscopic FP (Lorente de Nó, 1947; Nunez and Srinivasan, 2006; Herreras et al., 2022, 2023). Accordingly, although the relationship between FPs and spike activity in receiving populations is blurred, FPs can provide a window into upstream population dynamics (Elul, 1971; Herreras, 2016). To better understand this transformation, the present study asked whether FP features emerge from population-level signal integration or also reflect properties of elementary synaptic events.

Two features make micro-to-macro scaling particularly non-trivial: changes in source geometry and current cancellation. First, FPs are strongly shaped by how cellular morphology and population architecture determine which currents contribute to the extracellular field (Varona et al., 2000; López-Aguado et al., 2002; Fernández-Ruiz et al., 2013; Reimann et al., 2013; Gulyás et al., 2016; Ness et al., 2016; Telenczuk et al., 2020a; Herreras et al., 2023). Indeed, the joint spatial configuration of active sources can change with population activity, altering the extent and shape of the generating ensemble over time. Second, active (synaptic) and return currents (passive) can nearly cancel each other already within small tissue volumes, either within individual synaptic dipoles or among them, often resulting in little or no measurable potential (Lorente de Nó, 1947; Ahlfors et al. 2010; Herreras, 2016, Herreras et al., 2022, 2023). Consequently, neither the magnitude nor the temporal organization of population activity maps uniquely onto the magnitude or temporal structure of the resulting mesoscopic field at a given recording location, as demonstrated experimentally and reproduced computationally (Lindén et al., 2011; Fernández-Ruiz et al., 2013; Martín-Vázquez et al., 2016; Torres et al., 2019). The recorded FP therefore reflects the spatially filtered and potentially transformed expression of upstream activity, rather than providing a direct readout of that activity.

Large-scale models have explored how FP temporal dynamics emerge from population-level network organization (Robinson et al., 2003; Sanzleon et al., 2013; Kumar and Ma, 2023; Tesler et al., 2024; Meneghetti et al., 2025; Ness et al., 2025), but generally do not examine how microscopic current properties and the spatial geometry of the generating population jointly shape the resulting FP signals. At a different level, feed-forward modelling studies have shown that FP waveforms and conventional metrics such as amplitude and frequency can be strongly influenced by the morphoelectronic properties of receiving cells (Rall and Shepherd, 1968; Varona et al., 2000; López-Aguado et al., 2002; Brunel and Wang 2003; Geisler et al., 2005; Reimann et al., 2013; Martín-Vázquez et al., 2016; Ness et al., 2016).

A practical way to work around these difficulties without explicitly modelling source geometry, volume-conduction mixing, or time-varying current cancellation is to use spatially stable FP components obtained by independent component analysis (ICA) of experimental multisite recordings (Herreras et al., 2015). Unlike raw FPs, which mix contributions from simultaneously active sources, FP generators provide empirically derived representations of their net contributions. Their spatial profiles are defined by the ICA decomposition, while their time courses retain the fluctuations of the corresponding generator-specific contributions, reflecting upstream synaptic drive, including the net effects of current cancellation. This spatially collapsed representation thus retains the aggregate effects of source geometry and spatial averaging without requiring these factors to be reconstructed explicitly. FP generators have been obtained in rodents and humans (Fernández-Ruiz et al., 2012; Schomburg et al., 2014; Nasretdinov et al., 2023; Makarova et al., 2025), and some reflect individual synaptic pathways (Fernández-Ruiz et al., 2012; Benito et al., 2014, 2016). Although predominantly irregular (Hernández-Recio et al., 2023), they can be discriminated through combinations of spectral, nonlinear, and fractal metrics (Muñoz-Arnaiz et al., 2025). FP generators thus provide a controlled platform for testing whether mesoscopic features reflect the temporal organization of their inputs or also retain properties of elementary contributions.

We therefore asked whether the waveform properties of elementary synaptic-like extracellular contributions are reflected in the features used to characterize mesoscopic activity. To address this question, we developed a micro-to-mesoscopic approach to reconstruct experimentally obtained FP generators using unitary extracellular contributions (UECs) as proxies for elementary synaptic inputs. By keeping the temporal sequence of microscopic events identical across reconstructions while varying UEC waveform shape, we isolated the contribution of elementary waveform properties from that of temporal organization. UEC waveform influenced mesoscopic amplitude, spectral composition, and a broad range of nonlinear, fractal, and entropy-based features, with effects specific to the FP generator. Together, these findings indicate that properties of elementary extracellular contributions can leave detectable imprints on mesoscopic FP activity and may contribute to the physiological interpretation of commonly used FP metrics.

## METHODS

### Experimental FP-generators

FP generators were reused from linear FP recordings in previously published datasets from the rodent laboratory database. Detailed surgical, recording, animal-care, and ethical procedures are reported in those publications (Montero-Atalaya et al., 2023; Hernández-Recio et al., 2023). To enhance generalizability, we used the same subset of four FP generators from ten animals recorded under the same experimental protocols, selecting the four generators that were most consistently identified across different activity regimes.

Generators were obtained by ICA from linear depth recordings in the CA1/DG hippocampal region. Among the selected generators, two were pathway-specific—the CA3–CA1 Schaffer pathway and the lateral perforant pathway (LPP) input to the dentate gyrus—whereas the other two were defined by their spatial origin: the stratum lacunosum-moleculare (LM) generator and a remote (Rem) generator volume-conducted from cortical sources (Benito et al., 2014; 2016; Torres et al., 2019). These generators therefore provided spatially coherent FP sources spanning both pathway-defined and spatially defined activity patterns. In each animal, all generators were extracted from the same recording epochs. Long-duration epochs (51 ± 16 min; range: 18–74 min) were selected as templates to maximize sampling of the intrinsic temporal variability of each generator and minimize dependence on arbitrarily selected short episodes. Periods dominated by theta oscillations were excluded to avoid excessive weighting of highly coherent activity during ICA decomposition (Makarova et al., 2011). For quantitative analyses, the resulting epochs were segmented into consecutive 5-s windows, a duration previously shown to contain sufficient information to discriminate FP generators using their signal signatures (Muñoz-Arnaiz et al., 2025).

### Reference and alternative UEC waveforms

A single UEC waveform was used as a simplified representation of the effective extracellular contribution of an elementary synaptic input within the spatially collapsed representation of the generator. Importantly, a UEC does not represent an individual synapse or a fixed number of synapses. Rather, it represents an effective, spatially weighted contribution, collapsed into a single non-spatial measure. Sources at different distances and orientations contribute unequally to the extracellular field, and their individual contributions are therefore not equivalent. Accordingly, the number of UECs required to reproduce a given FP amplitude should be regarded as an indicative scaling parameter reflecting the effective density of active contributing sources, rather than as a direct estimate of the number or anatomical density of synapses (Lindén et al., 2011; Łęski et al., 2013; Reimann et al., 2013; Toharia et al., 2016; Hagen et al., 2017).

Six UEC waveforms were used across reconstructions: three biologically plausible alpha functions lasting 20 ms (alpha1, alpha2, and alpha3; Tau1 = 3, 9, and 0.1 ms, respectively; Tau2 = 1, 2, and 10 ms, respectively) and three artificial geometric waveforms lasting 10 ms (triangle, sine, and square; Fig. 2). One waveform from each category (alpha1 and sine) served as a reference UEC, providing two reference surrogates per trial. Each reference waveform was used to generate a surrogate event train, which was then convolved with all six UEC waveforms, yielding twelve reconstructions per trial. The two reference reconstructions were included in the analysis because they were also used to calculate feature differences relative to the original signal. This design allowed the effects of UEC waveform shape to be assessed independently of the temporal structure of the surrogate trains.

All UECs were normalized to the same peak amplitude, and the total discrete integral of each waveform relative to a zero baseline was calculated. A nominal amplitude of 0.5 nV was assigned to provide a plausible order-of-magnitude correspondence with the effective, spatially weighted extracellular contribution expected from synapses within a volume corresponding to 2 × 1 × 0.1 mm³ of CA1 stratum radiatum for the Schaffer input. This nominal amplitude was applied uniformly to all UECs across generators. It was intended as a reference for scaling the reconstructed signals, rather than as a direct estimate of unitary synaptic contributions, which depend strongly on morphoelectronic properties and source geometry (Ibarz et al., 2006; Lindén et al., 2011; Herreras et al., 2022). Variations in this nominal amplitude would change the absolute scale of the reconstructed signals but not the relative waveform-dependent effects examined in this study.

### Elaboration of surrogate event trains and generation of FP-generator reconstructions

Because each FP generator originates from a distinct neuronal population with its own endogenous input dynamics, natural input trains were not modeled; instead, surrogate event trains with a standardized temporal sequence were used to remove population-specific temporal organization as a source of variation. Two different surrogate trains were generated separately for each generator and animal using the two reference UECs (2 reference trains x 6 UECs x 4 FP generators x 10 animals = 480 reconstructions). UECs were assigned to time points at 1-ms intervals with a random jitter of ± 0.5 ms, producing temporal stacking of UECs around each selected instant, providing a controlled approximation of synchronous or near-synchronous activation of multiple inputs. This temporal stacking can resemble the concurrent activation of neuronal assemblies (Abeles et al., 1993; Yuste et al., 2024), whereby multiple synaptic inputs become concurrently active and contribute to the buildup of mesoscopic fields. At each time point, UECs were cumulatively added until their summed envelope reached the corresponding amplitude of the original FP-generator signal. Reconstructions therefore assumed linear superposition of elementary synaptic contributions and did not account for nonlinear interactions among convergent or overlapping inputs at individual neurons.

### Features explored

FP generators and their reconstructions were compared in terms of frequency content, using power spectral density (PSD) in predefined frequency bands: delta (0.5-3.5 Hz), theta (3.5-10 Hz), alpha-beta (10-30 Hz), gamma (30-100 Hz), and high-frequency (100-180 Hz). The micro-to-macro fidelity was assessed by comparing original and reconstructed signals through the linear cross-correlation (CC), phase-locking value (PLV), probability density histograms (PDH), and spectral coherence. Statistical significance of PLV and spectral coherence were assessed using surrogate tests (n = 1000). The time lag between the original and reconstructed signals at the CCmax value was also computed (TAUmax).

Additional features were calculated for each 5-s segment of the original and reconstructed generators. These included distributional and spectral statistics (signal skewness and kurtosis, and the mean, variance, skewness, and kurtosis of the spectrum and power spectral density), fractal and nonlinear measures (Higuchi FD, Katz FD, and correlation dimension), and complexity and information-theoretic measures (negentropy, spectral entropy, SVD entropy, permutation entropy, approximate entropy, sample entropy, WEntropy, and WEntropy–Rényi). Definitions and computational details of the less common features, together with their possible physiological interpretation, are provided in the Supplementary Methods.

### Statistics

Statistical analyses were conducted at the animal level (n = 10), with repeated observations within each animal treated as paired measurements. The four experimentally identified FP generators (Schaffer, LM, LPP, and Rem) were retained as separate within-animal conditions, and generator-dependent effects were assessed across animals.

Reconstruction fidelity was assessed using CCmax and TAUmax calculated between the original FP generators and their corresponding reconstructions. Differences across the six UEC waveforms and across the four FP generators were assessed using Friedman tests, with Kendall’s W reported as a measure of effect size. TAUmax was calculated relative to the alpha1 waveform. Phase- and frequency-domain fidelity was further assessed using probability density histograms (PDH), phase-locking value (PLV), and spectral coherence over 0–180 Hz. PLV and PDH were compared across frequency bands using Friedman tests, with Kendall’s W reported as a measure of effect size. Spectral coherence was used to assess frequency-dependent similarity between reconstructed and experimental signals, with statistical significance assessed using surrogate tests (n = 1000).

For signal amplitude and variability, reconstructed signals were analyzed in unfiltered (DC) and mean-centered (AC-like) modes. Differences in DC amplitude and AC variability among the four FP generators were assessed using Friedman tests. UEC-dependent effects were assessed using paired Wilcoxon signed-rank tests or Friedman tests, as appropriate, followed by post-hoc paired comparisons with Holm correction. For AC variability, additional analyses were performed after normalization by the area of each UEC waveform to determine whether waveform-dependent differences could be explained by differences in UEC area.

Spectral density was compared across the four FP generators and across the original signals and their six UEC-based reconstructions using Friedman tests, with Kendall’s W reported as a measure of effect size. The distribution of generator- and waveform-dependent effects across frequency bands was subsequently examined descriptively.

For each feature, generator, and UEC waveform, corresponding original and reconstructed 5-s segments were paired, and Cohen’s d was calculated for each pair of windows. Window-level effect sizes were then averaged within each animal to obtain a single animal-level effect size for each feature, generator, and UEC condition. These animal-level effect sizes were tested against zero using Wilcoxon signed-rank tests across the ten animals. The 21 features and six UEC conditions yielded 126 feature–UEC comparisons for each surrogate template, and p-values were corrected for multiple comparisons using the Benjamini–Hochberg false discovery rate (FDR) procedure.

Generator-dependent differences in standardized effect sizes were assessed using Friedman tests across the four generators. Post-hoc paired comparisons were performed only for feature–UEC combinations showing a significant omnibus generator effect, with correction for multiple comparisons using the Benjamini–Hochberg FDR procedure. The same analyses were repeated for surrogate event trains generated using the α1 and sine UEC waveforms to assess the reproducibility of UEC- and generator-dependent effects across surrogate templates.

All tests were two-sided, and statistical significance was set at p < 0.05 unless otherwise specified.

## RESULTS

The analysis pipeline is summarized in Figure 1. Four homologous, spatially stable FP generators were selected from ICA analysis of linear hippocampal recordings obtained from 10 animals by different experimenters (see Methods). These were: the Schaffer pathway (Sch), the lateral perforant pathway (LPP), the stratum lacunosum-moleculare (LM), and a remote volume-conducted component (Rem).These FP generators were baseline-adjusted to compensate for the artificial polarity reversal introduced by AC coupling, thereby representing each generator as a single-polarity contribution at the recording site (Martín-Vázquez et al., 2013). For each generator, surrogate microscopic event trains were generated by cumulatively adding UECs of a single waveform type until the amplitude of the experimental generator was reached. Two reference UEC waveforms, alpha1 and sine, representing the biologically plausible and geometrical waveform categories, respectively, were used to generate independent surrogate event trains. Each resulting event train was then used to generate six separate reconstructions, each using a single UEC waveform, comprising three biologically plausible and three geometrical shapes (Figure 2). Thus, the temporal sequence of events was kept identical across reconstructions derived from the same reference train, while only the UEC waveform was varied. The reconstructed signals were subjected to the same AC filtering, segmented into 5-s epochs, and characterized using 21 signal metrics.

**Figure 1.**
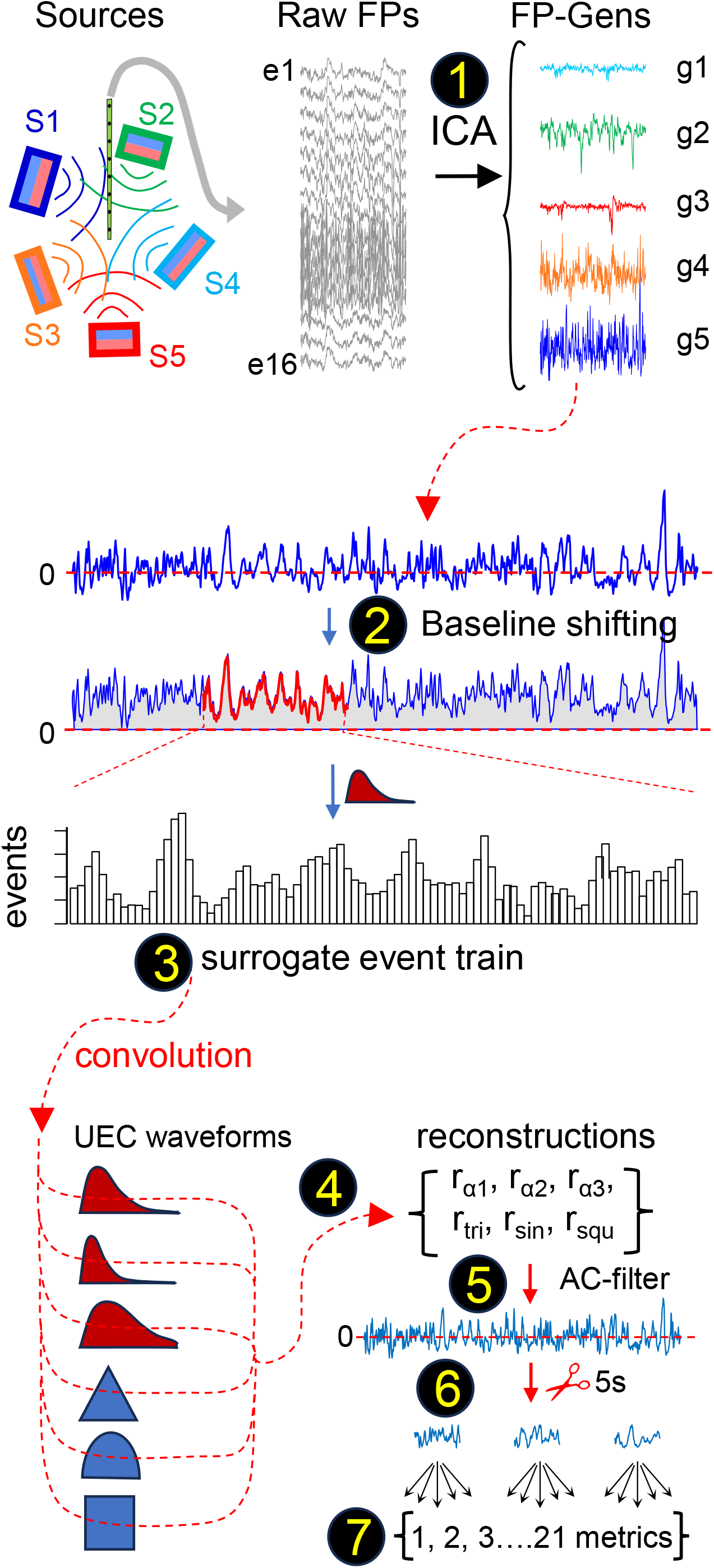
Reconstruction of field-potential generators from unitary extracellular contribution waveforms. Schematic of the micro-to-mesoscopic reconstruction procedure. (1) Raw field-potential (FP) recordings from multiple sources were decomposed by independent component analysis (ICA) into FP generators. (2) Each selected FP generator was shifted to a single polarity before reconstruction. (3) A surrogate event train was generated from the temporal profile of the FP generator. (4) The event train was convolved separately with six UEC waveforms (α1, α2, α3, triangular, sine and square) to obtain six reconstructions for each surrogate train and FP generator. (5) Reconstructions were AC-filtered to match the frequency range of the recorded FP signals. (6) The resulting signals were divided into 5-s epochs. (7) Twenty-one signal features were extracted from each epoch for comparison with the original FP generators.

**Figure 2.**
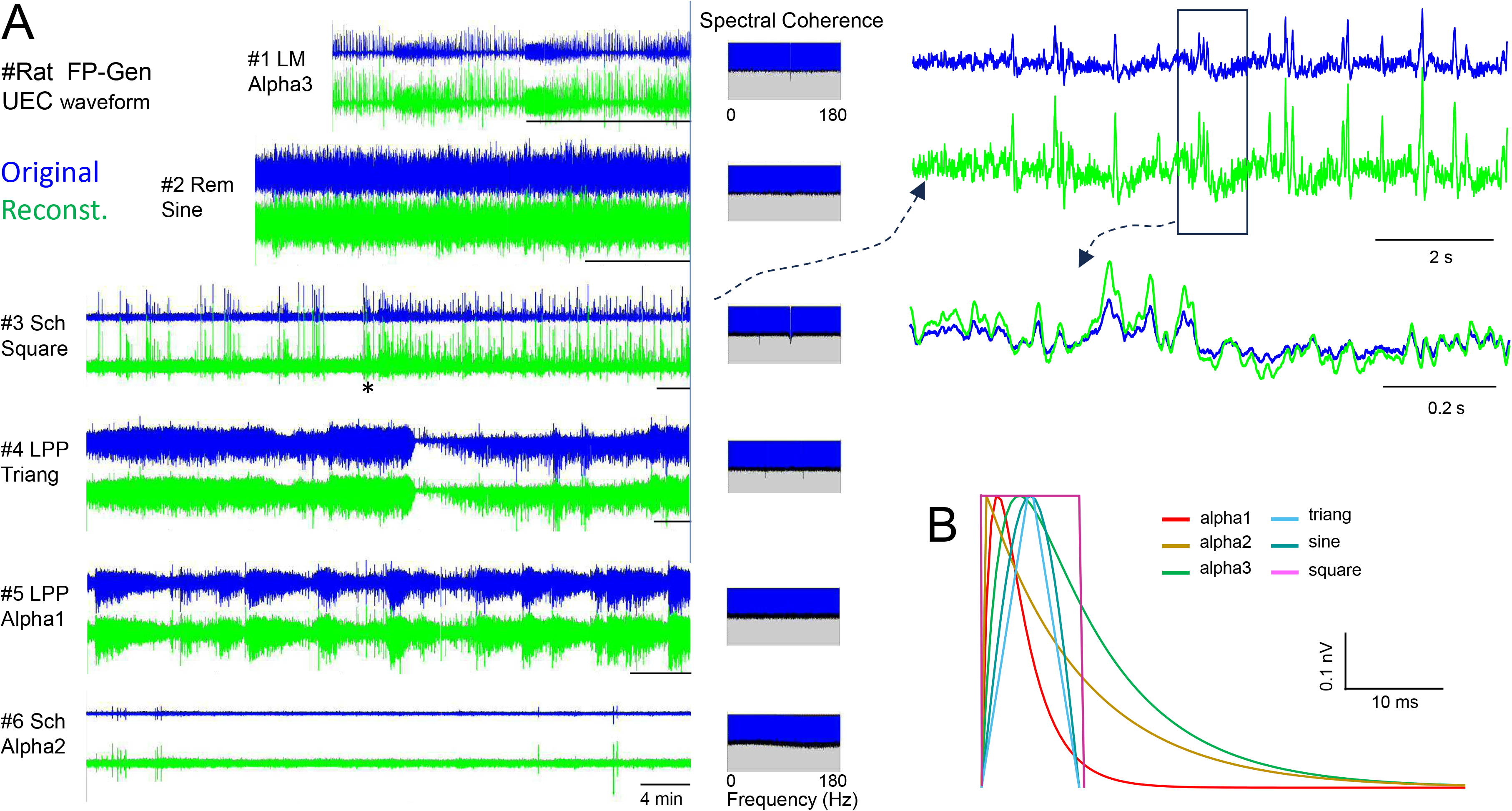
Reconstruction of field-potential generators using different UEC waveforms. (A) Examples of original FP-generator signals (blue) and their reconstructions (green) obtained using different UEC waveforms. Six examples are shown, combining different rats, FP generators (Sch, LPP, Rem and LM), and UEC waveforms (α1, α2, α3, triangular, sine and square). Left, extended signal segments; middle, spectral coherence between original and reconstructed signals from 0–180 Hz: blue color reflects the significant portion; right, expanded time segments illustrating the similarity between original and reconstructed signals. The six UEC waveforms used for reconstruction, normalized to the same peak amplitude to isolate the effects of waveform shape.

We first assessed whether the reconstruction procedure retained the main temporal and spectral characteristics of the original FP generators. We then examined how changing UEC waveform affected signal amplitude, spectral composition, and higher-order features, while keeping the surrogate event sequence unchanged. This allowed us to distinguish waveform-dependent effects from those arising from differences in temporal organization.

### High fidelity with high-frequency phase instability

The reconstruction showed high fidelity despite the length and temporal complexity of the templates, which displayed multiple visually distinct temporal patterns across FP generators. Figure 2 illustrates in six representative examples the close correspondence between the original templates and their reconstructions generated using different UEC waveforms (see waveforms in Fig. 2B). Consistent with this visual correspondence, maximum cross-correlation (CCmax) was high across all conditions, with values generally approaching unity (Fig. 3A). Nevertheless, CCmax showed small but systematic differences across UEC waveforms (Friedman χ²(5)=46.36, p<0.001, Kendall’s W=0.927) and generators (χ²(3)=23.47, p<0.001, W=0.782). TAUmax, measured relative to the reference α waveform, also differed systematically across UEC waveforms (χ²(5)=48.97, p<0.001, W=0.979), whereas no generator effect was detected (χ²(3)=5.18, p=0.159, W=0.173). The same qualitative waveform-dependent pattern was observed for alpha1- and sine-based surrogate trains, although the numerical TAUmax values differed between surrogate types. Interestingly, in both cases, the three geometric waveforms showed more similar TAUmax values to each other and greater separation from the alpha-function waveforms (Fig. 3B). Overall, these results confirm that the reconstructed signals retained high fidelity to the experimental FP generators, while preserving small waveform-dependent differences in temporal alignment.

**Figure 3.**
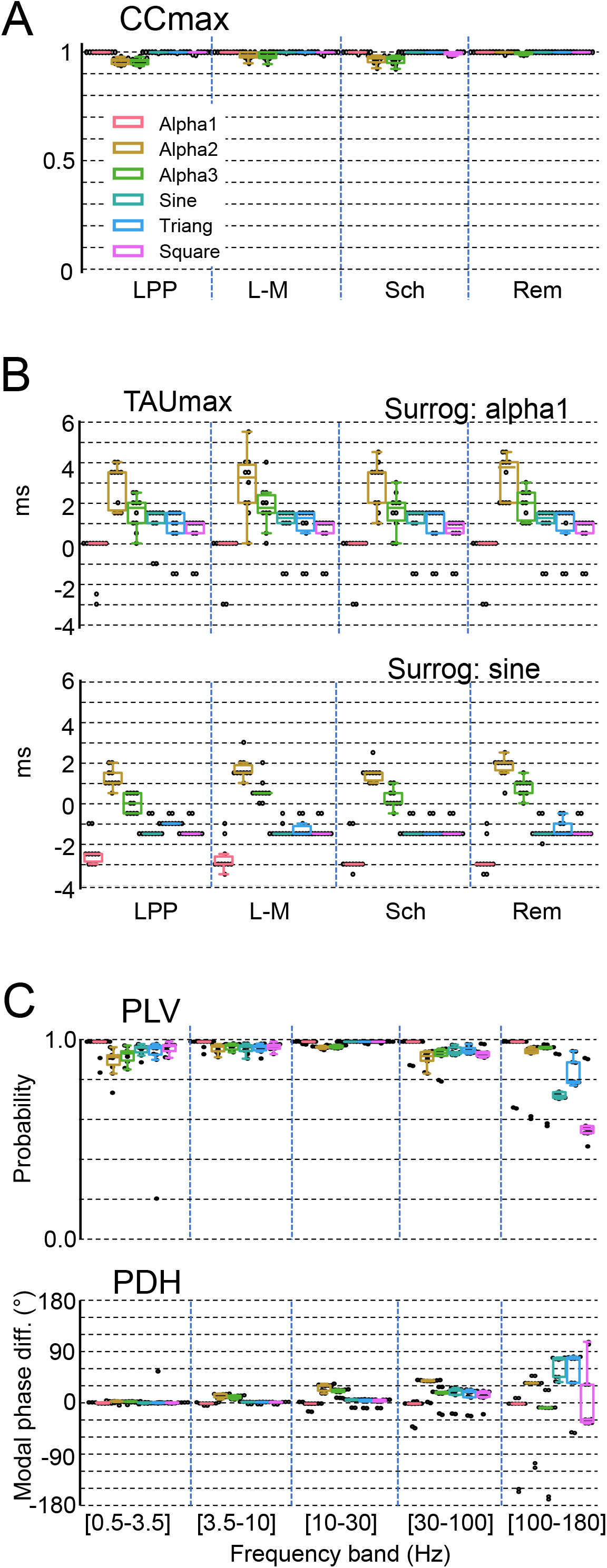
Temporal and phase correspondence between original and reconstructed FP generators. (A) Maximum cross-correlation (CCmax) between original and reconstructed FP generators for the four generators (LPP, L-M, Sch and Rem) and six UEC waveforms (α1, α2, α3, sine, triangular and square). (B) Time lag at maximum cross-correlation (TAUmax) for reconstructions obtained from alpha1- and sine-based surrogate event trains. (C) Phase-locking value (PLV; upper) and modal phase difference (PDH; lower) between original and reconstructed signals across five frequency bands. Box plots show the distributions across animals.

Phase-related measures revealed systematic differences across frequency bands and UEC waveforms in the reconstructed signals (Fig. 3C and Supplementary Fig. 1). Spectral coherence between reconstructed and experimental generators remained significant and close to unity across the analyzed frequency range (Fig. 2A). In contrast, the phase-locking values (PLV) varied significantly across frequency bands (χ²(4)=33.68, p<0.001, W=0.842), with a marked reduction at higher frequencies, indicating reduced phase consistency. This reduction was more pronounced for the sine, triangle, and square waveforms than for the three alpha-function waveforms. The phase difference histograms (PDH) also differed across frequency bands (χ²(4)=15.68, p=0.0035, W=0.392). These differences were barely discernible in the overall signal pattern when examined at a timescale of hundreds of milliseconds.

### UEC waveform impacts DC-amplitude and AC-variability of reconstructed FP-generators

The selected UECs were constrained to have the same peak amplitude, but not the same area under the curve. Consequently, accumulation of UECs along the surrogate event trains generated non-zero baselines in the reconstructed generators, upon which local fluctuations were superimposed (Fig. 4A). The mean amplitude of the unfiltered reconstructions, equivalent to DC recording, was proportional to the area of the UEC waveforms (Fig. 4B and Supplementary Fig. 2). This was expected because the same surrogate event train was used for all UEC waveforms within each animal and generator, making the UEC area the main scaling factor. Therefore, they primarily reflected differences already present in their original templates, rather than in the surrogate event timing. Reconstructed DC amplitude differed significantly among the four FP generators (Friedman test, χ²(3) = 12.60, p = 0.0056, Kendall’s W = 0.42). Post-hoc Wilcoxon signed-rank tests with Holm correction showed significantly larger amplitudes for the LPP generator than for the Sch and Rem generators (both p = 0.023), whereas no other pairwise comparisons reached significance. Across individuals, linear regression slopes were close to 1 for all reconstructions and surrogate trains. Population data showed greater dispersion, but the regression remained generator- and surrogate-train-specific (for LPP, LM, Sch, and Rem, R² values for the α1-based reconstructions were 0.78, 0.54, 0.82, and 0.50, respectively, and 0.72, 0.52, 0.84, and 0.46 for the sine-based reconstructions; Fig. 4C). Reconstructions generated from the sine reference train had consistently lower amplitudes than those generated from the α1 reference train (21% across all conditions; paired Wilcoxon signed-rank tests, p = 0.00195; Fig. 4A), matching the corresponding difference in the reference UEC amplitudes and further supporting a mainly multiplicative scaling effect rather than a generator-specific effect (Fig. 4B).

**Figure 4.**
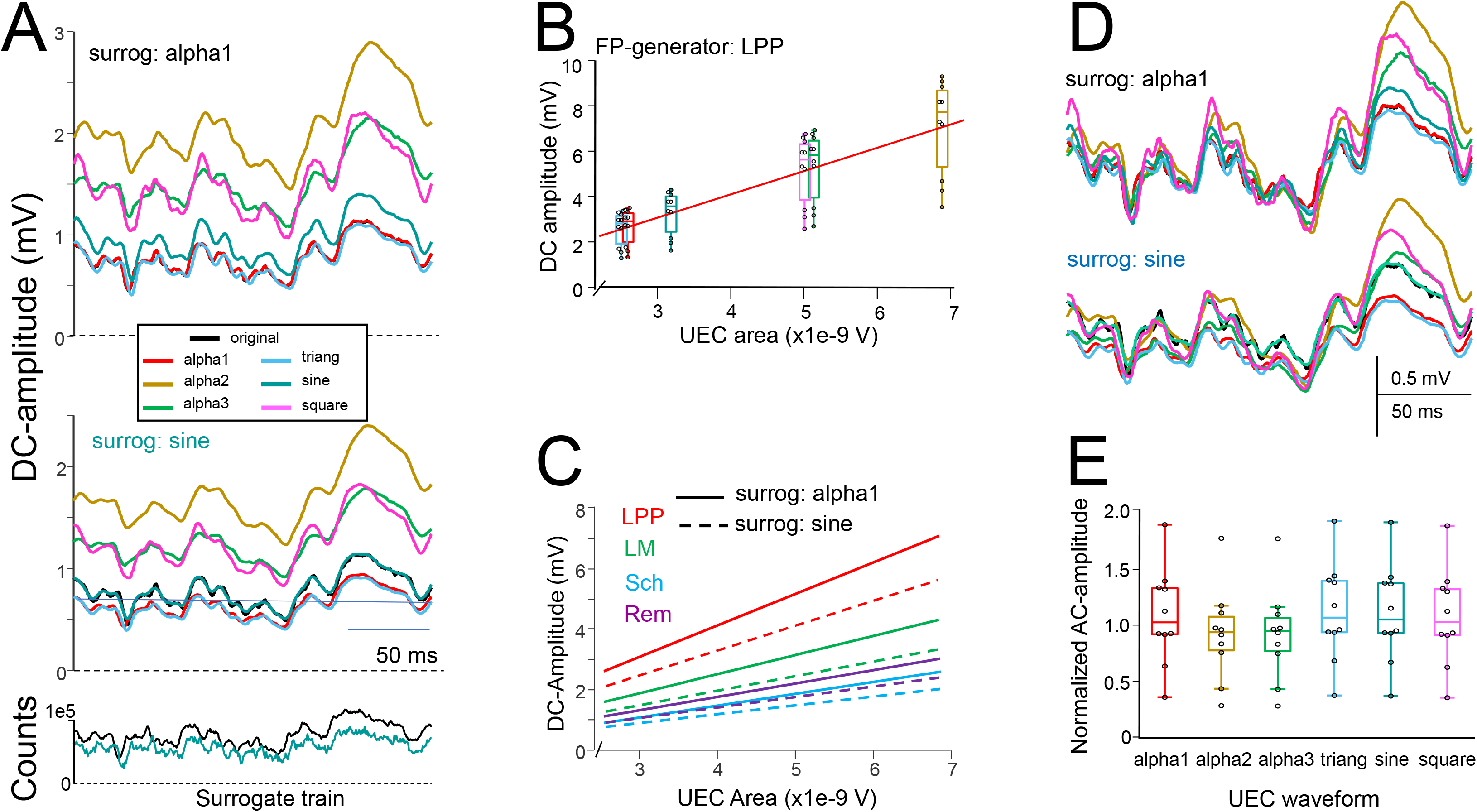
UEC waveform effects on DC amplitude and AC variability of reconstructed FP generators. (A) Examples of reconstructed FP-generator signals obtained from alpha1-(upper panel) and sine-based surrogate event trains (middle panel). The original generator is shown in black and reconstructions using the six UEC waveforms are shown in color. The corresponding surrogate event trains are shown below. (B) Relationship between UEC area and mean DC amplitude for an example LPP generator. Each box represents a reconstruction using one of the six UEC waveforms. (C) Linear relationships between UEC area and DC amplitude for the four FP generators, shown separately for Alpha1-(solid lines) and sine-based (dashed lines) surrogate event trains. (D) Expanded examples illustrating the effect of UEC waveform on the reconstructed signal for Alpha1- and sine-based surrogate trains. (E) Normalized AC amplitude of the reconstructions as a function of UEC waveform.

We next asked whether variability of the reconstructed signal could likewise be explained by differences in UEC area. Variability was quantified as the standard deviation of the reconstructed signal around its mean, thereby capturing fluctuations independently of the non-zero DC baseline. This is equivalent to standard AC-like recording mode. Visual inspection showed clear waveform-dependent differences, particularly for large-amplitude and shorter-duration waveforms (Fig. 4D). Generator identity significantly affected reconstructed variability (Friedman test, χ²(3) = 21.96, p < 0.001, Kendall’s W = 0.73). We then tested whether these differences, as well as the waveform-dependent effects, could be explained by the different UEC areas. Normalization by UEC area did not remove the dependence on waveform shape (Fig. 4E), which remained highly consistent across animals with the same ordering for all four generators (α3 < α2 < square < α1 < sine < triangle; Friedman tests, all χ²(5) = 50.00, p < 0.001, Kendall’s W = 1). All pairwise comparisons between waveforms remained significant after Holm correction. Thus, waveforms with identical peak amplitude but different temporal profiles generated markedly different levels of macroscopic variability even after accounting for their overall area. The same pattern was reproduced using an independent surrogate event train generated with the sine UEC (Supplementary Fig. 3).

### UEC waveform has a strong impact on spectral properties of the reconstructed signals

The reduced phase consistency observed at higher frequencies led us to examine whether these differences were reflected in the spectral properties of the reconstructed signals. The spectral density was strongly dependent on both generator identity (χ²(3) = 22.44, p < 0.001, Kendall’s W = 0.748) and UEC waveform (χ²(6) = 58.63, p < 0.001, Kendall’s W = 0.977) (Fig. 5). These effects were not uniformly distributed across the frequency spectrum. Generator-dependent differences were most consistently observed in the α, β, γ1, and γ2 bands, whereas effects in δ were restricted to the Schaffer generator and no significant generator-related interaction was detected in θ. Similarly, the effect of waveform varied across frequency bands, with the strongest and most consistent waveform-dependent effects occurring in the α, δ, and θ bands, whereas γ2 showed little evidence of a waveform-dependent effect. Thus, the main determinants of the reconstructed spectral density were generator identity and UEC waveform, with their effects being selectively expressed across frequency bands. This suggests that spectral differences between signals originating from different structures should not necessarily be attributed solely to differences in population dynamics, as intrinsic properties of the elementary contributions may also shape the resulting spectral content.

**Figure 5.**
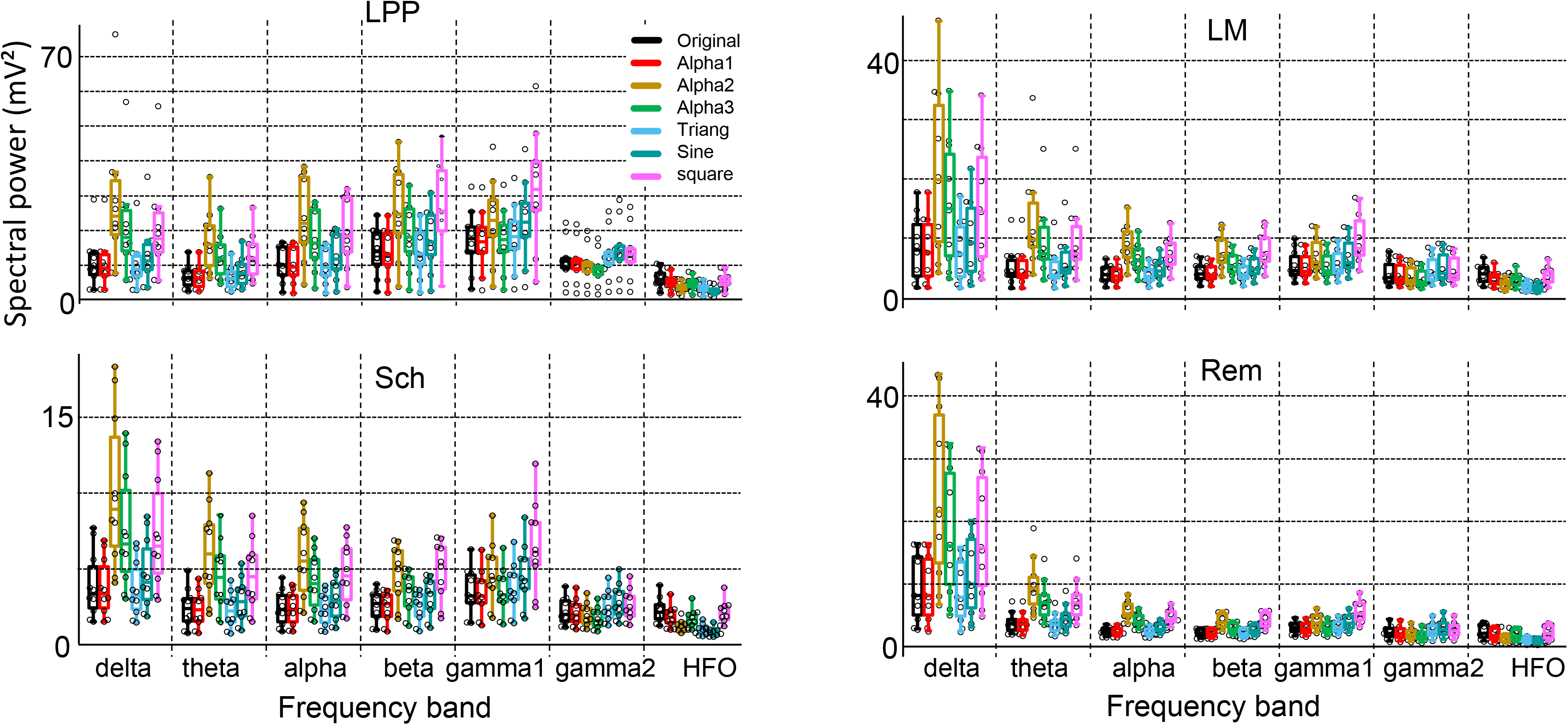
UEC waveform effects on the spectral power of FP-generator reconstructions. Spectral power across seven frequency bands (HFO: high-frequency oscillations) for the four FP generators. The original FP-generator signal (black) is compared with reconstructions obtained using the six UEC waveforms. Box plots show the distribution across animals for each generator, frequency band and waveform.

### UEC waveform effects on higher-order signal properties

UEC waveform affected a broad range of extracted signal features, with significant changes detected for 112 of the 126 feature–UEC combinations across the four generators (surr: alpha1) after FDR correction (Fig. 6 and Supplementary Figure 4). Significant effects were found across all feature classes, including distributional statistics, spectral and power spectral density measures, fractal/dimensional descriptors, and entropy- and complexity-based measures. Correlation dimension, Higuchi and Katz fractal dimensions, and several entropy measures were among the features most consistently affected by UEC waveform. Spectral and power spectral density descriptors were also frequently affected, indicating that waveform-dependent changes were not limited to the temporal statistics of the reconstructed signals.

**Figure 6.**
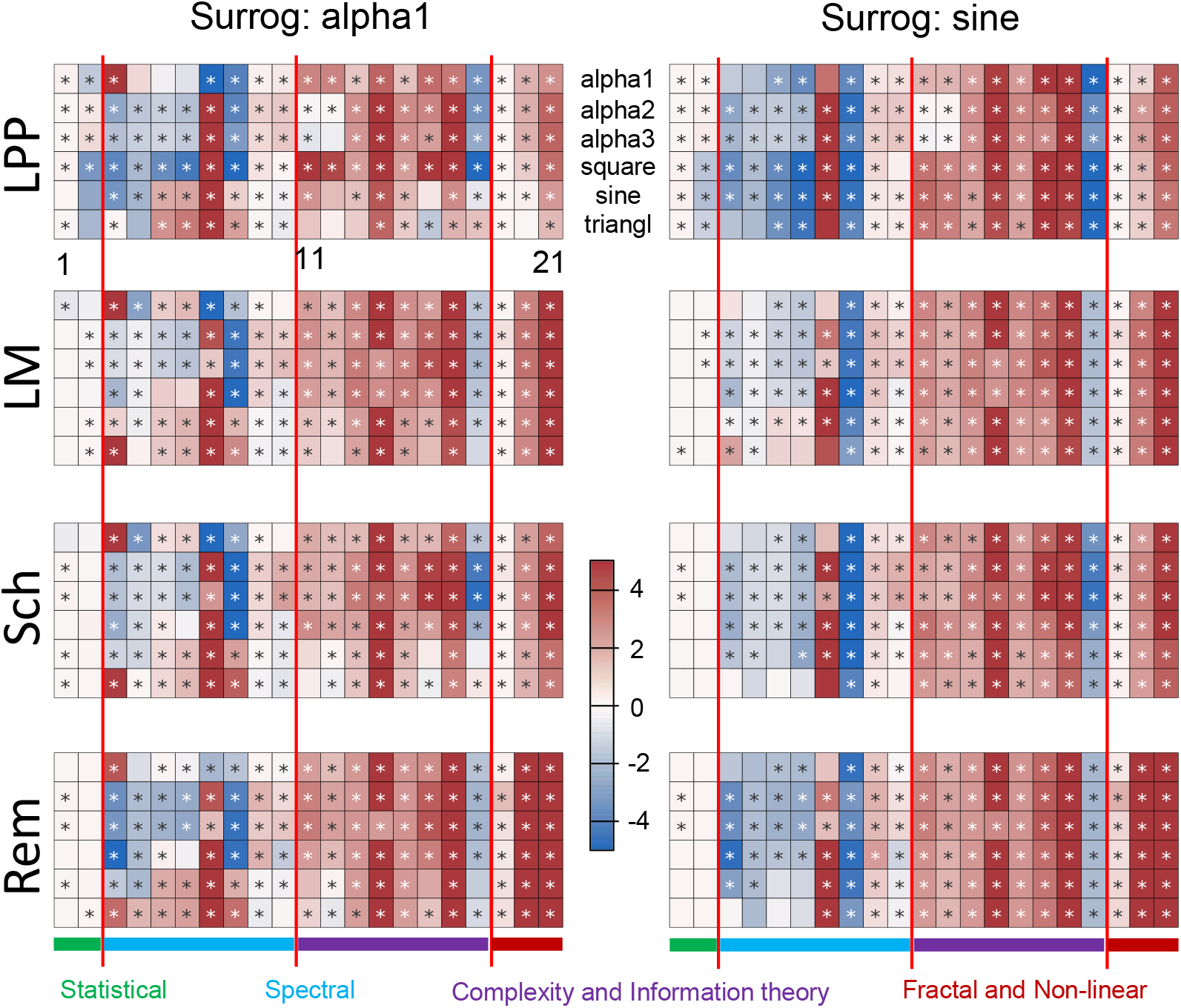
UEC waveform effects on extracted signal features. Heatmaps showing the standardized effect size of the difference between reconstructed and original FP-generator signals for 21 signal features. Results are shown separately for the four FP generators and for reconstructions based on Alpha1- and sine-based surrogate event trains. Rows within each heatmap correspond to the six UEC waveforms, and columns correspond to the 21 features grouped into statistical, spectral, complexity and information-theory, and fractal and nonlinear measures. Color indicates the direction and magnitude of the standardized effect size; asterisks indicate significant differences after FDR correction.

The magnitude of the effects differed among generators. Friedman tests showed significant differences in standardized effect size among GD, LM, Sch, and Rem for 86 of the 126 feature–UEC combinations after FDR correction (Fig. 7). These generator-dependent differences were found across all six UECs, with 12–16 significant feature–UEC combinations per UEC, and were particularly frequent for correlation dimension, Higuchi and Katz fractal dimensions, PSD variance and skewness, and spectral variance. Pairwise comparisons were then performed for the 86 feature–UEC combinations with a significant generator effect, yielding 516 generator-pair comparisons, of which 274 remained significant after FDR correction. Thus, the magnitude of the UEC-related effects differed among generator types. Complete effect-size results for all UEC waveform–feature comparisons are provided in Supplementary Data 1 and 2.

**Figure 7.**
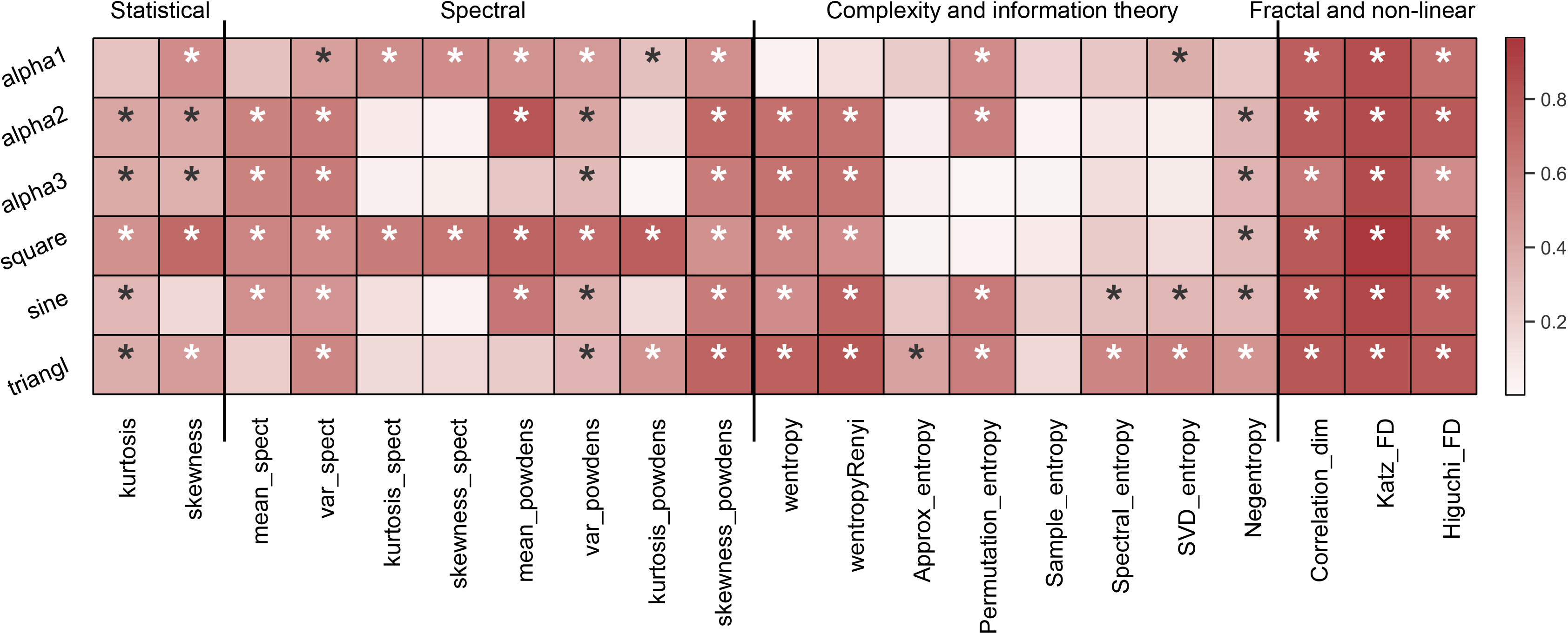
Generator-dependent effects of UEC waveform across signal features. Heatmap showing the Friedman test effect size (Kendall’s W) for differences in standardized effect size among the four FP generators, calculated across the ten animals for each UEC waveform and signal feature. Rows correspond to the six UEC waveforms, and columns to the 21 features grouped into statistical, spectral, complexity and information-theory, and fractal and nonlinear measures. Color intensity indicates the magnitude of Kendall’s W. Asterisks indicate significant generator-dependent differences after Benjamini–Hochberg FDR correction.

We next examined whether these effects were also observed when the surrogate template was changed from alpha1 to sine. Significant effects were observed for 412 of the 504 feature–UEC–generator combinations in both analyses, indicating substantial overlap between the two surrogate templates. Generator-dependent differences were also observed in both analyses for 69 feature–UEC combinations. However, 36 combinations showed significant effects only with alpha1 and 31 only with sine, while generator-dependent differences were exclusive to alpha1 or sine for 17 and 11 combinations, respectively. Thus, most UEC effects and generator-dependent differences were reproduced across surrogate templates, although some feature-specific effects depended on the surrogate used.

## DISCUSION

The generic goal of this study was to explore how microscopic current properties are transformed when scaling to mesoscopic FPs. Despite the large number of UECs required to generate measurable FP amplitudes, their waveform properties influenced DC amplitude, AC variability, spectral properties, and nonlinear and complexity measures, with effects that depended on the FP generator.

### Making tractable a multilevel scaling phenomenon

The microscopic-to-mesoscopic transformation of synaptic activity into FPs involves a large number of geometrical and functional factors operating at different organizational scales. Previous modeling approaches have addressed this complexity by using intermediate representations of neuronal and synaptic sources, with varying degrees of biological detail and experimental constraint (Brunel, 2000; Bojak et al., 2010, 2011; Sanz-Leon et al., 2013; Hendrickson et al., 2015; Markram et al., 2015; Breakspear, 2017; Telenczuk et al., 2020; Li et al., 2021; Tesler et al., 2022). Such approaches make multilevel modeling feasible, but necessarily restrict which microscopic factors can be examined independently and how directly their contributions can be related to macroscopic signals.

Synaptic geometry, dendritic location, electrotonic properties, and voltage-dependent channels all affect the extracellular mixing and cancellation of synaptic and return currents (Lindén et al., 2011; Reimann et al., 2013; Ness et al., 2016; Herreras et al., 2023). Their interactions make individual contributions difficult to isolate. We therefore adopted a reconstruction strategy that bypasses the need to explicitly model three major sources of uncertainty: the unknown geometry of microscopic sources, the mixing of signals from different synaptic pathways in raw FPs, and the time-dependent cancellation of extracellular currents (Herreras, 2016). The use of FP generators rather than raw, volume-conducted FPs as templates is central to this strategy. ICA provides spatially coherent representations of individual FP generators, which can often be associated with activity in identified pathways (Herreras et al., 2015). It also separates remote volume-conducted contributions, which are often overlooked despite dominating FP recordings in many brain structures (Nunez and Srinivasan, 2006; Tenke and Kaiser, 2012; Martín-Vázquez et al., 2016; Torres et al., 2019). This reduces the confounding effects of simultaneously active sources, whose overlapping waveforms result in raw FP waveforms that do not faithfully copy either source individually (Makarova et al., 2011; Torres et al., 2019; Herreras et al., 2022), and helped establish that the predominance of irregular activity reflects the intrinsic dynamics of upstream neuronal populations rather than mutual distortion during source mixing (Hernández-Recio et al., 2023; see also Dehghani et al., 2010). Finally, their experimental origin ensures that fluctuations reflect the time-varying fraction of extracellular currents surviving cancellation, meaning that measured AC amplitudes should not be taken at face value even apart from the fact that they exclude the DC component,.

Using FP generators, however, required a spatially abstract proxy for synaptic currents: the UEC. Its assumed amplitude limits the relation between the number of UECs and the underlying number of synaptic sources. Nearby sources contribute strongly, whereas distant sources contribute little individually but may be numerous (Lindén et al., 2011; Łęski et al., 2013; Reimann et al., 2013). The UEC therefore represents a spatially weighted contribution collapsed into a single non-spatial measure. The number of UECs should consequently be regarded as an indicative scaling parameter rather than a direct estimate of synaptic density or source number. Whereas mean-field models necessarily rely on assumptions about numerous architectural and spatial factors, often without explicitly accounting for their implications, the use of FP generators allows the present UEC design to bypass the need to model these factors explicitly.

Several factors that normally shape FP fluctuations were not incorporated in the present reconstruction. Most importantly, UEC recruitment timing was neutralized by using the same surrogate event train across waveforms within each generator. This allowed us to isolate the contribution of UEC waveform properties, but does not address how recruitment timing, synchrony, or other forms of temporal organization interact with microscopic current properties. Such interactions necessarily influence spectral and other macroscopic properties and warrant separate examination. The present results therefore describe one component of the microscopic-to-mesoscopic transformation rather than a complete model of FP generation.

### UEC waveform impacts visible and non-visible features of the FPs

High fidelity of visible FP features might be expected, but the preservation of generator-specific patterns in nonlinear features is less obvious. Despite using the same surrogate event train across UEC waveforms, generator-specific patterns remained detectable across the feature maps. This supports the view of FP generators as stable representations of distinct source populations (Herreras et al., 2015) and indicates that less visible nonlinear features can also distinguish generators, as previously shown by Muñoz-Arnaiz et al. (2025).

FP analysis has traditionally focused on properties visible in the time or frequency domains, such as waveform configuration, oscillations, amplitude, and spectral composition. However, much of the structure of an irregular FP is not visually apparent. Entropy, fractal, and nonlinear descriptors can capture statistical and temporal organization within apparently noisy fluctuations (Stam, 2005; Muñoz-Arnaiz et al., 2025). Their sensitivity to UEC waveform properties suggests that these features may contain information arising from the microscopic-to-mesoscopic transformation, rather than being merely abstract mathematical descriptors. Irregular FP fluctuations may therefore contain structured information about the microscopic-to-mesoscopic transformation that is not apparent from discrete events or visible oscillations.

### DC-level and AC variability, two sides of the same phenomenon impacted by microscopic waveforms

The persistence of UEC waveform effects after scaling to mesoscopic FPs might seem unexpected given the large number of events involved, with mean counts on the order of tens of thousands. Unlike previous modeling studies that varied the temporal correlation of synaptic inputs while keeping their number and location constant (Lindén et al., 2011; Reimann et al., 2013), we used temporally aligned UEC events because the actual timing of microscopic recruitment is unknown. Although this alignment may resemble the co-activation of synapses during assembly firing (Harris et al., 2003; Yuste et al., 2024), it may also have favored the preservation of UEC waveform effects, which was not a design objective.

Conversely, fully asynchronous activation would be expected to yield a more stable DC-like potential envelope, depending on the average rate and spatial balance of the underlying currents. Sustained DC potentials are now rarely considered, although they may be more prominent than generally assumed. While they are often associated with pathological conditions (Herreras and Makarova, 2020), they were the subject of extensive research in past decades and were also linked to cognitive states and task performance (Kovac et al., 2018). In our reconstructions, cumulative addition of UECs produced non-zero, DC-like baselines whose amplitude increased with UEC energy. Because conventional recordings are typically AC-coupled, this component cannot be fully captured experimentally, although DC shifts have been reported for well-characterized FP activities such as the theta rhythm (Brankack et al., 1993) and for so-called ultraslow oscillations (Cooper et al., 2025).

Interestingly, we also observed a proportional increase in AC variability with mean DC amplitude, in a UEC-dependent manner, although its impact on the reconstruction differed according to the amplitude and duration of the macroscopic waveform. This relationship persisted after UEC energy normalization, suggesting a role for the distribution of energy within the UEC waveform itself. Given that the temporal organization of microscopic events would be expected a priori to have a stronger influence on visible FP properties than the waveform of individual events, the persistence of UEC-dependent effects on AC variability is notable. This interaction between UEC energy, DC offset, and macroscopic waveform properties deserves further investigation.

### Toward understanding irregular activity with unconventional features

Efforts are increasingly being made to characterize the predominantly irregular activity of the brain using approaches that go beyond conventional signal descriptors (Donoghue et al., 2020). Our results add to this perspective by showing that the properties of elementary extracellular contributions affect different features of irregular FP activity to different extents. Changes in UEC waveform did not produce a uniform change in signal complexity or irregularity; their effects differed across feature classes and depended on the FP generator. This confirms that these descriptors are embedded in the real fluctuations of pathway-specific FP generators that are preserved across animals, and suggests that microscopic waveform properties can differentially contribute to the features used to distinguish them. If distinct source populations were characterized by different elementary synaptic current waveforms, due for example to differences in receptor kinetics, cellular properties, dendritic filtering, or other source-specific factors, these microscopic differences could contribute to distinct macroscopic feature profiles even under similar recruitment patterns. Thus, part of the feature-specific signature of an FP generator may originate from the properties of its elementary currents rather than exclusively from differences in population activity.

Different descriptors may capture distinct aspects of the microscopic-to-mesoscopic transformation. Fractal dimensions, such as Higuchi and Katz, characterize aspects of signal geometry and temporal structure, whereas spectral entropy reflects the distribution of energy across frequencies (Higuchi, 1988; Katz, 1988; Inouye et al., 1991). Temporal entropy measures, including permutation, approximate, and sample entropy, quantify different aspects of regularity and temporal predictability, while wavelet-based entropy describes energy distribution across temporal scales (Pincus, 1991; Richman and Moorman, 2000; Bandt and Pompe, 2002; Rosso et al., 2006). These measures capture complementary properties of FP fluctuations, although their sensitivity to UEC waveform changes cannot yet be assigned to specific physiological mechanisms.

This may also explain the heterogeneous effects observed across the 21 measures. This heterogeneity may arise because changes in UEC waveform can simultaneously affect fine-scale roughness, spectral distribution, temporal regularity, and multiscale organization, which different descriptors capture to varying degrees. The frequent effects observed for fractal and nonlinear measures suggest that some consequences of UEC waveform changes may be expressed more strongly in the organization of signal fluctuations than in conventional measures of amplitude or spectral power. Although these measures cannot yet be assigned to specific physiological mechanisms, our results suggest that different feature families may capture distinct aspects of the microscopic-to-mesoscopic transformation (Lau et al., 2022; Averna et al., 2023).

### Concluding remark

Mesoscopic field potentials are not shaped only by when populations are active, but also by what their elementary currents contribute to the signal. Our results show that microscopic waveform properties can leave detectable signatures across amplitude, spectral, and nonlinear features despite the massive integration of elementary events. This suggests that some properties of mesoscopic activity may reflect not only population organization, but also the microscopic currents from which it emerges.

## Data availability

The data supporting the findings of this study are available in the main text and Supplementary Information. The original electrophysiological recordings are available from the corresponding author upon reasonable request.

## Acknowledgements

We thank former lab members Sara Hernández-Recio and Marta Montero-Atalaya, who performed some of the experiments from which the FP recordings used in this study were obtained.

## Funding

This work was supported by grants PID2022-137801NB-I00 and CPP2023-010818 from the Spanish Ministries MICIN and MCIU, and EU2020/2094-IASOMM24006 from the CSIC excellence programs. RMA is financed by a PhD fellowship PIPF-2023/SAL-GL-30443 from the Comunidad Autónoma de Madrid.

## Author Contributions

Conceptualization: RMA, OH; Methodology: RMA, ARF, VAM; Investigation: RMA, FA, JM, OH; Visualization RMA, ARF, FA, JM, OH; Funding acquisition: OH; Writing original draft OH; Writing-review & editing: RMA, AFR, FA, JM, OH.

## Competing Interests

The authors declare no competing interests.

## Supplementary Methods

### Metrics definitions

The following expressions describe the mathematical formulation of the measures computed in this work. They are expressed for a given time series, with sampling frequency *F_s_* and sample mean. Unless stated otherwise, *log* denotes the natural logarithm.

### Temporal and spectral measures

#### Skewness

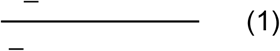

The skewness measures the asymmetry of a distribution. Zero skewness corresponds to a symmetric distribution. For an FP-type time series, it has no simple interpretation as it may be influenced by several parameters concerning individual waveforms or their grouping.

#### Kurtosis

Non-excess kurtosis (equal to 3 for a Gaussian variable):

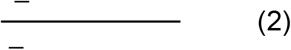

Kurtosis estimates the amount of data in the tails of a probability distribution related to the standard one. Gaussian noise has a kurtosis of 3. Due to the Central Limit Theorem (Montgomery and Runger, 2014), a large mixture of independent sources tends to have a standard distribution (i.e., becomes similar to noise). For an FP-type time series, kurtosis quantifies how many waves (actually, time points) stand out from the regular signal, i.e., are outliers. ICA maximizes deviations of the distributions from the normal one. Thus, the kurtosis of FPs is usually closer to 3 than that of FP generators that capture pathway-specific events.

### Spectral measures

Power spectra are commonly used to pinpoint the presence and relative power of rhythmic oscillations in an FP. Note however that any single FP wave already contains power in a wide power band. Brief periods of oscillatory activity could indeed be present and may strongly determine spectral properties in some segments. However, a given frequency band (e.g., gamma) is not sufficient discriminant feature as it may appear in several generators (Torres et al., 2019; López-Madrona et al., 2020; Hernández-Recio et al., 2023). Here we used global descriptors of the spectral domain (mean, variance, kurtosis and skewness) that may better account for the contribution of non-periodic activity. The power on specific standard bands was also estimated but not included in the group of measures due to low informativeness.

For the spectral mean, variance, kurtosis and skewness, the discrete Fourier transform is calculated:

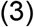

with associated frequencies.

#### Power spectral density (PSD)

For a wide-sense stationary process, the PSD describes the distribution of power across the frequency components of *x_n_* and is the Fourier transform of the autocorrelation function (Wiener–Khinchin theorem):

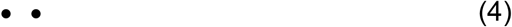

For an ergodic process, *R_xx_* can be computed as a time average,. From a finite record we use the periodogram estimate

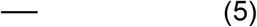

### Measures of entropy

The entropy of a random variable can be interpreted as the amount of information provided by the observation of a variable. The more random and unpredictable the variable is, the greater its entropy (Rényi, 1961).

For a discrete probability distribution, Shannon entropy is

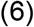

and Rényi entropy of order *α≠1* is

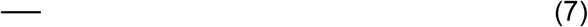

#### Wavelet entropies

A maximal overlap discrete wavelet transform (MODWT) with the Symlet 4 (sym4), with the decomposition level determined automatically from the signal length. Let *W_j_[n]* be the wavelet coefficients at scale *j*. The relative energy at each level is

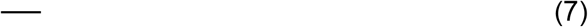

The Shannon (*H_S_*) and Rényi (H_α_) wavelet entropies are computed from with the formulas above, using α=2.

#### Spectral entropy

Let *P(F_j_)* be the PSD estimated with Welch’s method and

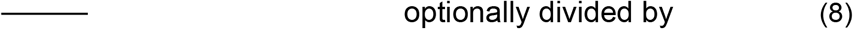

where *N_f_* is the number of frequency bins.

#### SVD entropy

Using an embedding of dimension 3 and delay 1, define

for *m = 1,2,3,* each of length *N-2*, and the matrix.

Let *M = UΣV^T^* be its singular value decomposition with singular values σ_1_, σ_2_, σ_3_. Then

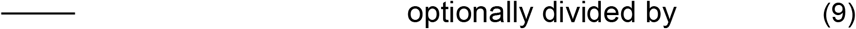

#### Permutation entropy

Form the vectors *v_n_ = (x_n_, x_n+1_, x_n+2_)*, *n=1,…N-2,* and map each to its ordinal pattern *π ɛ S_3_*. With *p(π)* the relative frequency of pattern *π* among the *N-2* vectors,

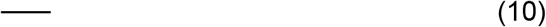

#### Approximate and Sample Entropies

A detailed discussion of both measures can be found in Delgado-Bonal and Marshak (2019).

### Other complexity measures

#### Negentropy

For a random variable *X* with differential entropy *H(X)*, negentropy is

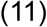

where *Xg* is a Gaussian variable with the same variance as *X*.

#### Correlation dimension D_2_

The correlation dimension (CD) of a set of points is a measure of its dimensionality. In general, CD describes the geometrical and dynamic complexity of a data set. However, at present, still there is room for studying its relation to FPs and biophysical interpretation of the obtained results (Mormann et al., 2005; Makarov et al., 2023). CD was initially created to estimate dimensionality of sets of points whose dimension is not an integer number.

Then, it has been extended to analysis of time series using Taken’s theorem (Sauer et al., 1991; Takens, 1981). We reported that the CD of FP generators follow closely each one’s temporal fluctuation (see Makarov et al., 2023 for procedural details).

The series is embedded in with delay, with *M = N-(m-1)Ƭ*. The correlation integral is

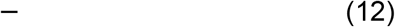

where ø is the Heaviside function and *N* is the number of pairs *(i,j)* with *j-1>w*. The correlation dimension is

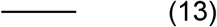

In practice, *D_2_* is estimated as the slope of *logC(r)* versus *logr*, with m = 3, Ƭ = 1.

#### Higuchi dimension D_H_

For an integer scale *k ≥ 1* and an initial point *m = 1,…,k*, consider the subseries and let. Its normalized length is

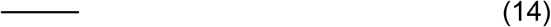

Then, *D_H_* is computed as:

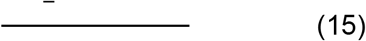

#### Katz dimension D_K_

Assuming a unit time step between samples,

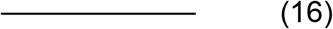

where *d* is the maximum distance from the first point and *L*:

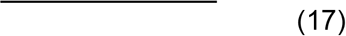

## Supplementary Figures

**Supplementary Figure 1.**
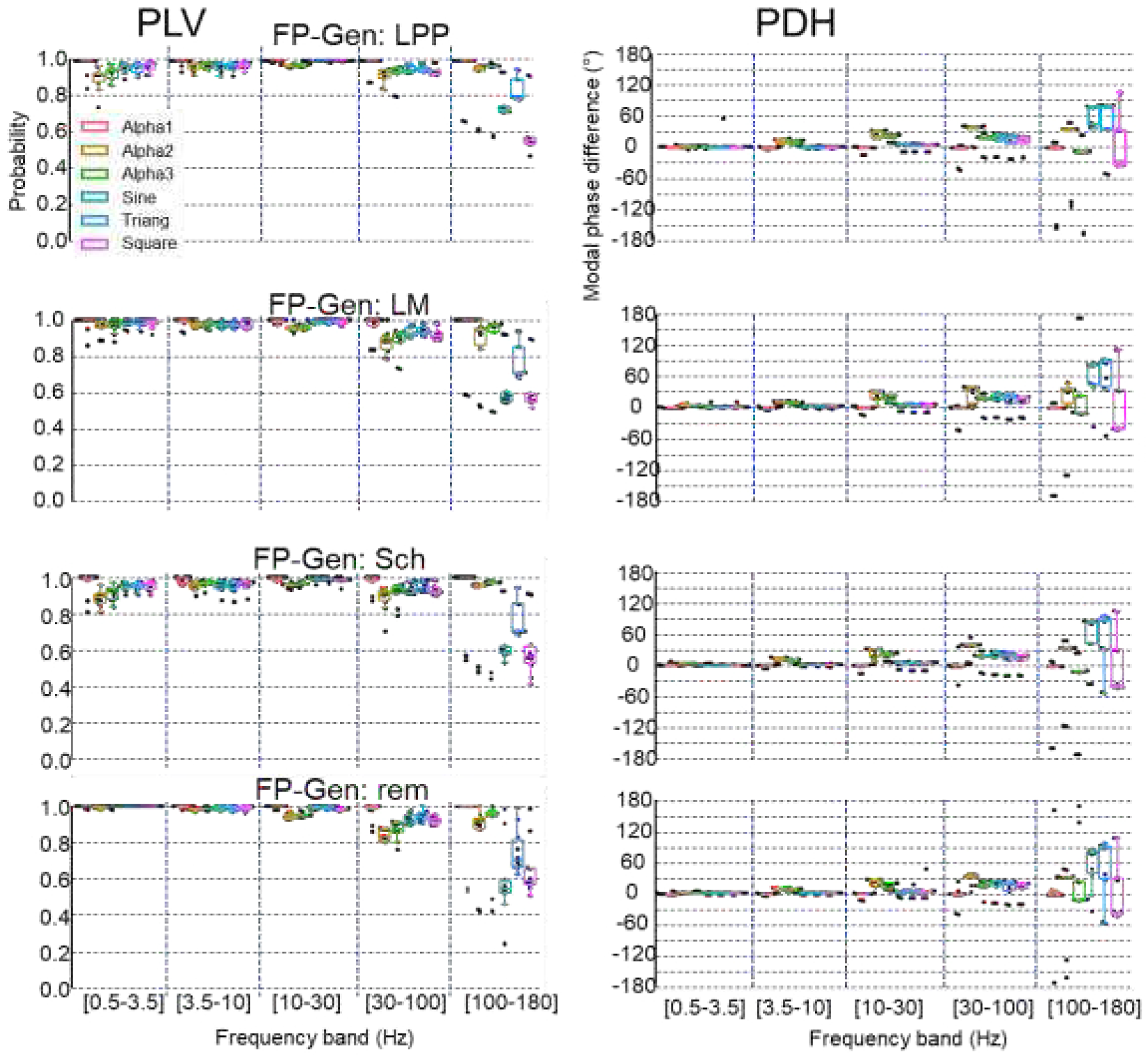
Phase correspondence between original and reconstructed FP generators. Phase-locking value (PLV) and modal phase difference (PDH) between original and reconstructed FP-generator signals across frequency bands. Results are shown for the four FP generators and the six UEC waveforms, using α1- and sine-based surrogate event trains.

**Supplementary Figure 2.**
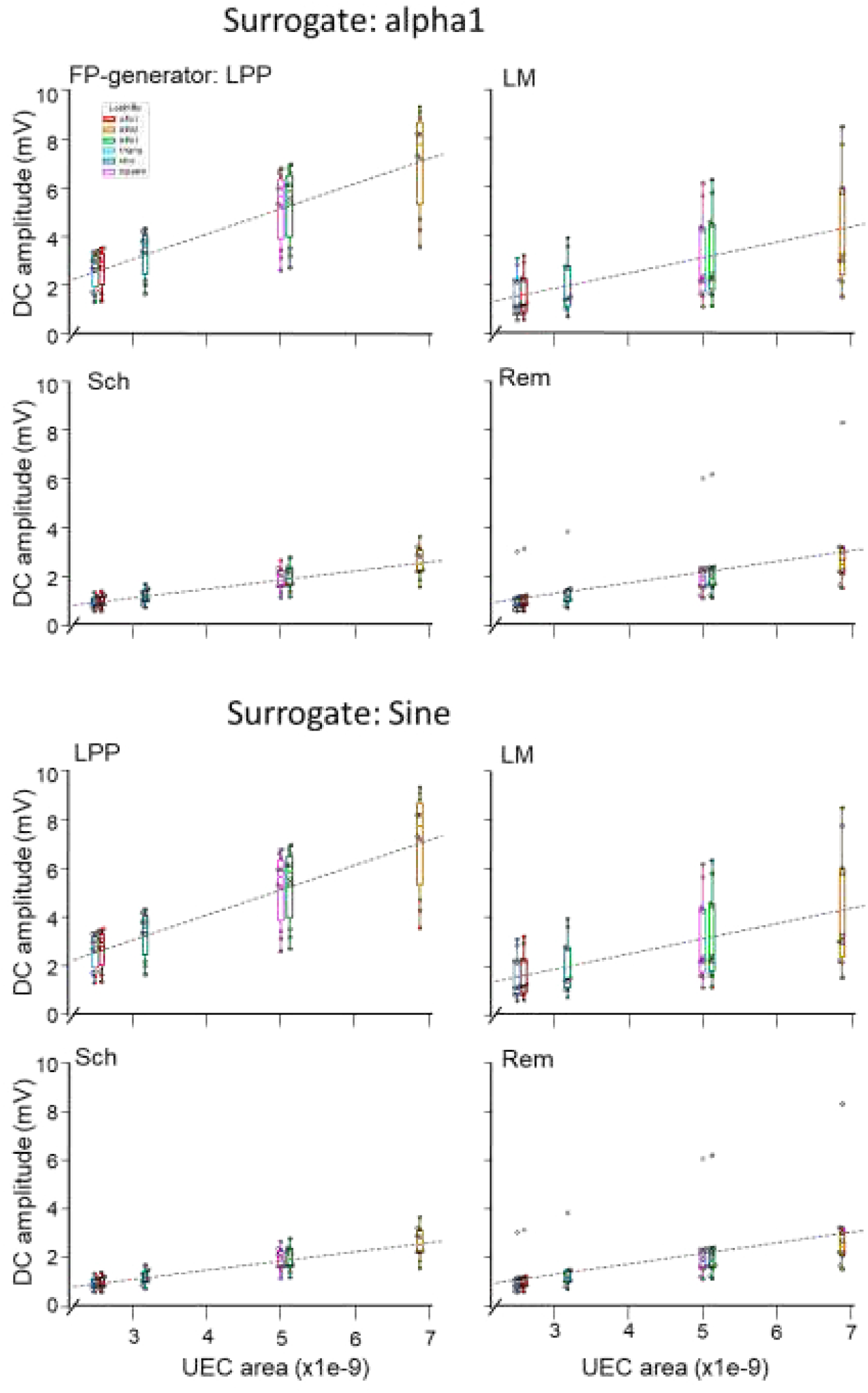
Relationship between UEC area and DC amplitude across FP generators. Relationship between UEC area and mean DC amplitude of the reconstructed FP generators for the four generators. Results are shown separately for alpha1- and sine-based surrogate event trains and for the six UEC waveforms. Each point represents a reconstruction, with linear fits shown for each FP generator. DC amplitude was measured from the unfiltered reconstructed signals.

**Supplementary Figure 3.**
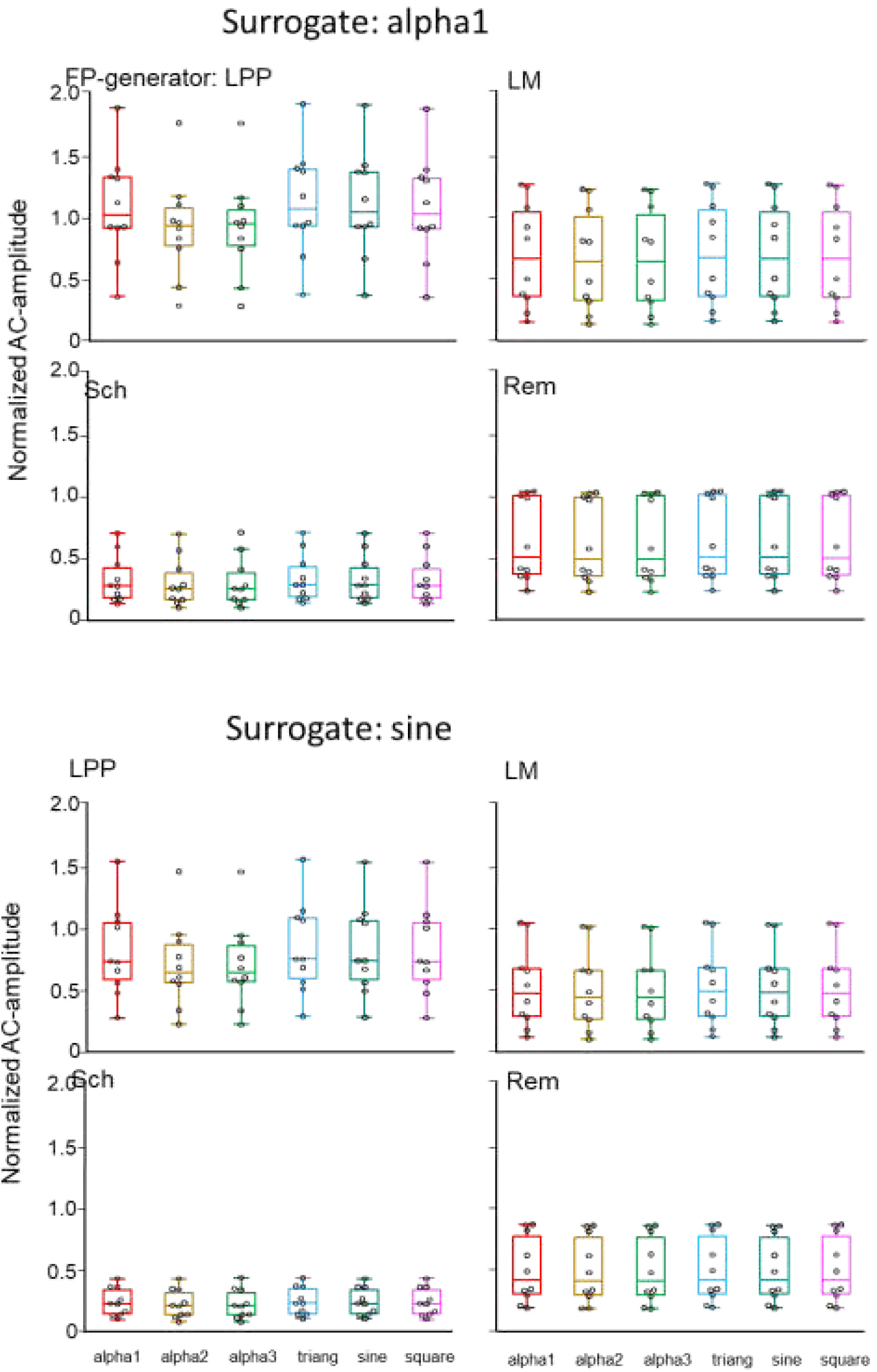
UEC waveform effects on AC amplitude across FP generators. Normalized AC amplitude of reconstructed FP generators for the four generators and six UEC waveforms. Results are shown for alpha1- and sine-based surrogate event trains. AC amplitude was measured after AC filtering and normalized to the corresponding original FP-generator signal.

**Supplementary Figure 4.**
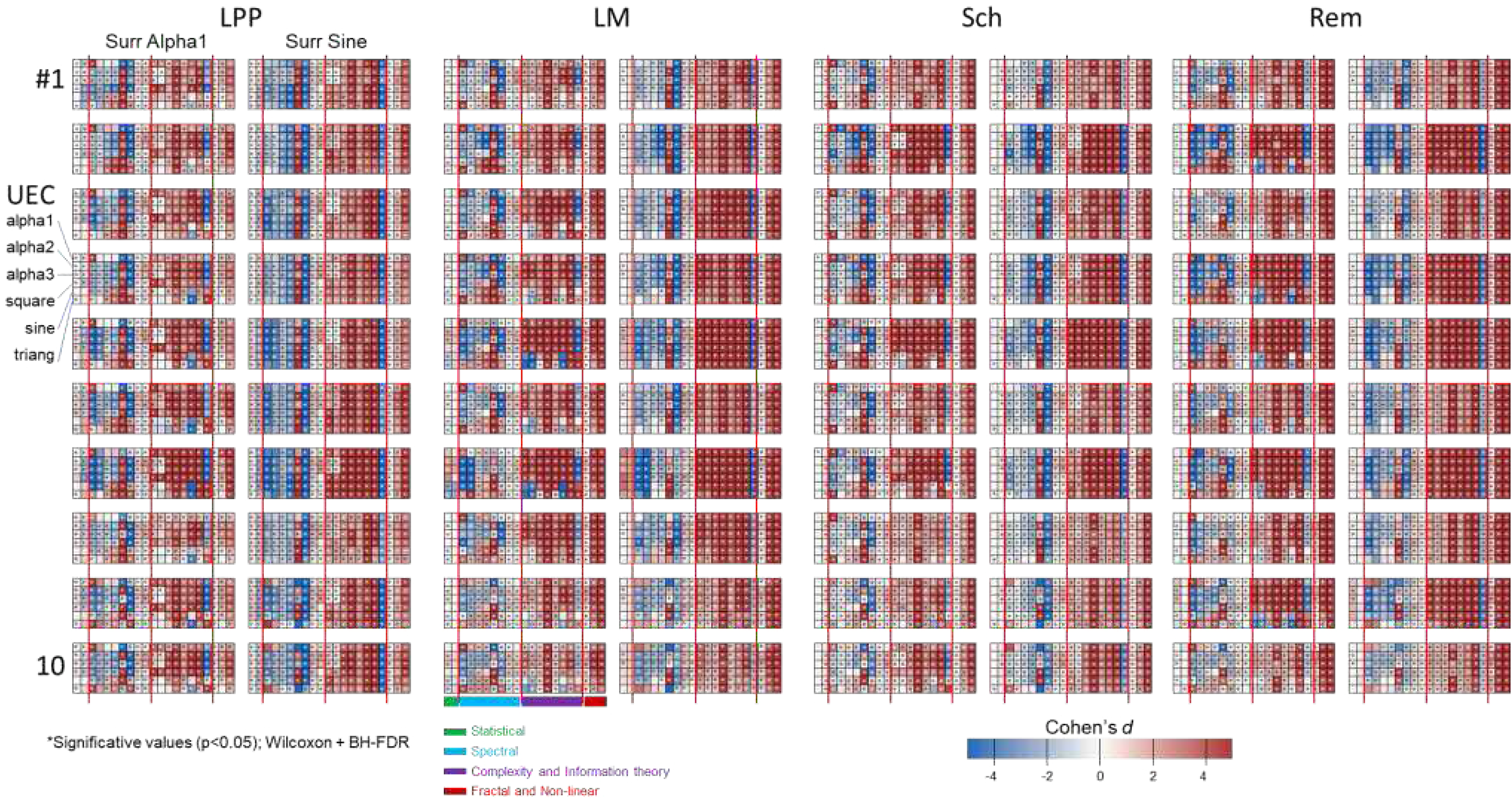
Effect sizes of UEC waveform changes across signal features and animals. Cohen’s d for the difference between reconstructed and original FP-generator signals across the 21 signal features. Results are shown for the four FP generators, the six UEC waveforms, and alpha1- and sine-based surrogate event trains. Features are grouped into statistical, spectral, complexity and information-theory, and fractal and nonlinear measures. Asterisks indicate significant differences after Wilcoxon tests with Benjamini–Hochberg FDR correction. Positive values indicate larger feature values in the reconstructed than in the original signals, whereas negative values indicate the opposite.

**Supplementary Data 1**

Effect sizes for all UEC waveform–feature comparisons in α1-based surrogate reconstructions. The dataset includes Cohen’s d values for paired comparisons between reconstructed and original field-potential generators, together with the corresponding statistical test results and multiple-comparison corrections. The complete dataset is provided as a separate file (cohensd_10n_alpha1.csv).

**Supplementary Data 2**

Effect sizes for all UEC waveform–feature comparisons in sine-based surrogate reconstructions. See the accompanying α1-based dataset for details. The complete dataset is provided as a separate file (cohensd_10n_sine.csv).

## Notes

### Competing Interest Statement

The authors have declared no competing interest.

## REFERENCES

Abeles, M., Bergman, H., Margalit, E., Vaadia, E. (1993). Spatiotemporal firing patterns in the frontal cortex of behaving monkeys. J Neurophysiol, 70, 1629–1638. doi:10.1152/jn.1993.70.4.1629.

Ahlfors, S.P., Han, J., Lin, F.H., Witzel, T., Belliveau, J.W., Hämäläinen, M.S., Halgren, E. (2010). Cancellation of EEG and MEG signals generated by extended and distributed sources. Hum Brain Mapp, 31, 140–149. 10.1002/hbm.20851

Averna, A., Coelli, S., Ferrara, R., Cerutti, S., Priori, A., Bianchi, A.M. (2023). Entropy and fractal analysis of brain-related neurophysiological signals in Alzheimer’s and Parkinson’s disease. J Neural Eng. 20, 5. doi:10.1088/1741-2552/acf8fa.

Bandt, C., Pompe, B. (2002). Permutation entropy: A natural complexity measure for time series. Phys Rev Letters, 88, 174102. 10.1103/PhysRevLett.88.174102.

Benito, N., Fernández-Ruiz, A., Makarov, V.A., Makarova, J., Korovaichuk, A., Herreras, O. (2014). Spatial modules of coherent activity in pathway-specific LFPs in the hippocampus reflect topology and different modes of presynaptic synchronization. Cereb Cortex, 24, 1738–1752. doi: 10.1093/cercor/bht022.

Benito, N., Martín-Vázquez, G., Makarova, J., Makarov, V.A., Herreras, O. (2016). The right hippocampus leads the bilateral integration of gamma-parsed lateralized information. eLife, 5, e16658. doi: 10.1093/cercor/bht022.

Bojak, I., Oostendorp, T.F., Reid, A.T., Kötter, R. (2010). Connecting mean field models of neural activity to EEG and fMRI data. Brain Topography, 23, 139–149. 10.1007/s10548-010-0140-3.

Bojak, I., Oostendorp, T.F., Reid, A.T., Kötter, R. (2011). Towards a model-based integration of co-registered electroencephalography/ functional magnetic resonance imaging data with realistic neural population meshes. Philos Trans A Math Phys Eng Sci, 369, 3785–3801. 10.1098/rsta.2011.0080.

Brankack, J., Stewart, M., Fox, S.E. (1993). Current source density analysis of the hippocampal theta rhythm: associated sustained potentials and candidate synaptic generators. Brain Res, 615, 310–327. 10.1016/0006-8993(93)90043-m.

Breakspear, M. (2017). Dynamic models of large-scale brain activity. Nat Neurosci, 20, 340–352. doi: 10.1038/nn.4497.

Brunel, N., Wang, X.J. (2003). What determines the frequency of fast network oscillations with irregular neural discharges? I. Synaptic dynamics and excitation-inhibition balance. J Neurophysiol, 90, 415–430. doi: 10.1152/jn.01095.2002.

Brunel, N. (2000). Dynamics of sparsely connected networks of excitatory and inhibitory spiking neurons. J Comput Neurosci, 8, 183–208. 10.1023/A:1008925309027.

Bullock, T.H., Mcclune, M.C, Enrigh, J.T (2003) Are the electroencephalograms mainly rhythmic? Assessment of periodicity in wide-band time series. Neuroscience, 121, 233–252. 10.1016/s0306-45220300208-2.

Buzsáki, G., Anastassiou, C.A., Koch, C. (2012). The origin of extracellular fields and currents--EEG, ECoG, LFP and spikes. Nat Rev Neurosci, 13, 407–420. doi: 10.1038/nrn3241.

Cooper, C., Parthier, D., Sibille, J., Tukker, J.J., Tritsch, N., Schmitz, D. (2025). Ultraslow serotonin oscillations in the hippocampus delineate substates across NREM and waking. Elife, 13, RP101105. doi: 10.7554/eLife.101105.

Dehghani, N., Cash, S.S., Rossetti, A.O., Chen, C.C., Halgren, E. (2010). Magnetoencephalography demonstrates multiple asynchronous generators during human sleep spindles. J Neurophysiol, 104, 179–188. 10.1152/jn.00198.2010.

Donoghue, T., Haller, M., Peterson, E.J., Varma, P., Sebastian, P., Gao, R., Noto, T., Lara, A.H., Wallis, J.D., Knight, R.T., Shestyuk, A., Voytek, B. (2020). Parameterizing neural power spectra into periodic and aperiodic components. Nat Neurosci, 23, 1655–1665. doi:10.1038/s41593-020-00744-x.

Eckhorn, R. (1994). Oscillatory and non-oscillatory synchronizations in the visual cortex and their possible roles in associations of visual features. Prog Brain Res, 102, 405–426. 10.1016/S0079-61230860556-7.

Elul, R. (1971). The genesis of the EEG. Int Rev Neurobiol, 15, 228–272.

Fernández-Ruiz, A., Makarov, V.A., Benito, N., Herreras, O. (2012). Schaffer-specific local field potentials reflect discrete excitatory events at gamma-frequency that may fire postsynaptic hippocampal CA1 units. J Neurosci, 32, 5165–5176. doi: 10.1523/jneurosci.4499-11.2012.

Fernández-Ruiz, A., Muñoz, S., Sancho, M., Makarova, J., Makarov, V.A., Herreras, O. (2013). Cytoarchitectonic and dynamic origins of giant positive LFPs in the Dentate Gyrus. J Neurosci, 33, 15518–15532. doi:10.1523/JNEUROSCI.0338-13.2013.

Geisler, C., Brunel, N., Wang, X.J. (2005). Contributions of intrinsic membrane dynamics to fast network oscillations with irregular neuronal discharges. J Neurophysiol, 94, 4344–4361. doi: 10.1152/jn.00510.2004.

Gulyás, A.I., Freund, T.F., Káli, S. (2016). The effects of realistic synaptic distribution and 3d geometry on signal integration and extracellular field generation of hippocampal pyramidal cells and inhibitory neurons. Front Neural Circuits, 10, 88. doi: 10.3389/fncir.2016.00088.

Hagen, E., Fossum, J.C., Pettersen, K.H., Alonso, J.M., Swadlow, H.A., Einevoll, G.T. (2017). Focal local field potential signature of the single-axon monosynaptic thalamocortical connection. J Neurosci, 37, 5123–5143. doi: 10.1523/JNEUROSCI.2715-16.2017.

Harris, K.D., Csicsvari, J., Hirase, H., Dragoi, G., Buzsáki, G. (2003). Organization of cell assemblies in the hippocampus. Nature, 424, 552–556. doi: 10.1038/nature01834.

He, B.J., Zempel, J.M., Snyder, A.Z., Raichle, M.E. (2010). The temporal structures and functional significance of scale-free brain activity. Neuron, 66, 353–369. 10.1016/j.neuron.2010.04.020.

Hendrickson, P.J., Yu, G.J., Song, D., Berger, T.W. (2015). A million-plus neuron model of the hippocampal dentate gyrus: critical role for topography in determining spatiotemporal network dynamics. IEEE Trans Biomed Eng, 63, 199–209. doi: 10.1109/TBME.2015.2445771.

Hernández-Recio, S., Muñoz-Arnaiz, R., López-Madrona, V., Makarova, J., Herreras, O. (2023). Uncorrelated bilateral cortical input becomes timed across hippocampal segments for long waves whereas gamma waves are largely ipsilateral. Front Cell Neurosci, 17, 1217081. 10.3389/fncel.2023.1217081.

Herreras, O., Makarova, J., Makarov, V.A. (2015). New uses for LFPs: pathway-specific threads obtained through spatial discrimination. Neuroscience, 310, 486–503. 10.1016/j.neuroscience.2015.09.054.

Herreras, O. (2016). Local field potentials: Myths and misunderstandings. Front Neural Circuits, 10, 101. 10.3389/fncir.2016.00101.

Herreras, O., Makarova, J. (2020). Mechanisms of the negative potential associated with Leao’s spreading depolarization: A history of brain electrogenesis. JCBFM, 40, 1934–1952. 10.1177/0271678X20935998

Herreras, O., Torres, D., Martín-Vázquez, G., Hernández-Recio, S., López-Madrona, V.J., Benito, N., Makarov, V.A., Makarova, J. (2022). Site-dependent shaping of field potential waveforms. Cereb Cortex, 33, 3636–3650. 10.1093/cercor/bhac297.

Herreras, O., Torres, D., Makarov, V.A., Makarova, J. (2023). Theoretical considerations and supporting evidence for the primary role of source geometry on field potential amplitude and spatial extent. Front Cell Neurosci, 17, 1129097. 10.3389/fncel.2023.1129097.

Higuchi, T. (1988). Approach to an irregular time series on the basis of the fractal theory. Physica D: Nonlinear Phenomena, 31, 277–283. 10.1016/0167-2789(88)90081-4.

Ibarz, J.M., Makarova, J., Herreras, O. (2006). Relation of apical dendritic spikes to output decision in CA1 pyramidal cells during synchronous activation: a computational study. Eur J Neurosci, 23, 1219–1233. doi:10.1111/j.1460-9568.2006.04615.x.

Inouye, T., Shinosaki, K., Sakamoto, H., Toi, S., Ukai, S., Iyama, A., Katsuda, Y., Hirano, M. (1991). Quantification of EEG irregularity by use of the entropy of the power spectrum. Electroenceph Clin Neurophysiol, 79, 204–210. 10.1016/0013-4694(91)90138-T.

Katz, M.J. (1988). Fractals and the analysis of waveforms Computers in Biology and Medicine 18, 145–156. doi: 10.1016/0010-4825(88)90041-8

Kovac, S., Speckmann, E.J., Gorji, A. (2018). Uncensored EEG: The role of DC potentials in neurobiology of the brain Prog Neurobiol 165–167, 51-65. doi:10.1016/j.pneurobio.2018.02.001.

Kumar, G., Ma, C.H.E. (2023). Toward a cerebello-thalamo-cortical computational model of spinocerebellar ataxia. Neural Netw, 162, 541–556. doi: 10.1016/j.neunet.2023.01.045.

Lau, Z.J., Pham, T., Chen, S.H.A., Makowski, D. (2022). Brain entropy, fractal dimensions and predictability: A review of complexity measures for EEG in healthy and neuropsychiatric populations. Eur J Neurosci, 56, 5047–5069. doi: 10.1111/ejn.15800

Łęski, S., Lindén, H., Tetzlaff, T., Pettersen, K.H., Einevoll, G.T. (2013). Frequency dependence of signal power and spatial reach of the local field potential. PLoS Comput Biol, 9(7), e1003137. doi:10.1371/journal.pcbi.1003137.

Li, G., Liu, Y., Zheng, Y., Wu, Y., Li, D., Liang, X., Chen, Y., Cui, Y., Yap, P.T., Qiu, S., Zhang, H., Shen, D. (2021). Multiscale neural modeling of resting-state fMRI reveals executive-limbic malfunction as a core mechanism in major depressive disorder. NeuroImage: Clinical, 31, 102758. 10.1016/j.nicl.2021.102758.

Lindén, H., Tetzlaff, T., Potjans, T.C., Pettersen, K.H., Grün, S., Diesmann, M., Einevoll, G.T. (2011). Modeling the spatial reach of the LFP. Neuron, 72, 859–872. doi:10.1016/j.neuron.2011.11.006.

López-Aguado, L., Ibarz, J.M., Varona, P., Herreras, O. (2002). Structural inhomogeneities differentially modulate action currents and population spikes initiated in the axon or dendrites. J Neurophysiol, 88, 2809–2820. 10.1152/jn.00183.2002

Lorente de Nó, R. (1947). Analysis of the distribution of the action currents of nerve in volume conductors. Stud Rockefeller Inst Med Res Repr, 132, 384–477.

Makarov, V.A., Makarova, J., Herreras, O. (2010). Disentanglement of local field potential sources by independent component analysis. J Comput Neurosci, 29, 445–457. doi:10.1007/s10827-009-0206-y

Makarova, J., Ibarz, J.M., Makarov, V.A., Benito, N., Herreras, O. (2011). Parallel readout of pathway-specific inputs to laminated brain structures. Front Syst Neurosci, 5, 77. doi: 10.3389/fnsys.2011.00077

Makarova, J., Toledano, R., Blázquez-Llorca, L., Sánchez-Herráez, E., Gil-Nagel, A., DeFelipe, J., Herreras, O. (2025). Intracranial voltage profiles from untangled human deep sources reveal multisource composition and source allocation bias. J Neurosci, 45(1), e0695242024. doi: 10.1523/JNEUROSCI.0695-24.2024.

Markram, H., Muller, E., Ramaswamy, S., Reimann, M.W., Abdellah, M., Sanchez, C.A., Ailamaki, A., Alonso-Nanclares, L., Antille, N., Arsever, S., Kahou, G.A., Berger, T.K., Bilgili, A., Buncic, N., Chalimourda, A., Chindemi, G., Courcol, J.D., Delalondre, F., Delattre, V., et al (2015) Reconstruction and simulation of neocortical microcircuitry. Cell, 163, 456–492. 10.1016/j.cell.2015.09.029.

Martín-Vázquez, G., Makarova, J., Makarov, V.A., Herreras, O. (2013) Determining the true polarity and amplitude of synaptic currents underlying gamma oscillations of local field potentials. PLoS ONE, 8(9), e75499. doi:10.1371/journal.pone.0075499.

Martín-Vázquez, G., Benito, N., Makarov, V.A., Herreras, O., Makarova, J. (2016). Diversity of LFPs activated in different target regions by a common CA3 input Cereb Cortex, 26, 4082–4100. doi:10.1093/cercor/bhv211.

Meneghetti, N., Rimehaug, A.E., Einevoll, G.T., Mazzoni, A., Ness, T.V. (2025). Kernel-based LFP estimation in detailed large-scale spiking network model of mouse visual cortex. Neural Netw, 192, 107866. doi: 10.1016/j.neunet.2025.107866.

Montero-Atalaya, M., Expósito, S., Muñoz-Arnaiz, R., Makarova, J., Bartolomé, B., Martín, E., Moreno-Arribas, M.V., Herreras, O. (2023). A dietary polyphenol metabolite alters CA1 excitability ex vivo and mildly affects cortico-hippocampal field potential generators in anesthetized animals. Cereb Cortex, 33, 10411–10425. doi: 10.1093/cercor/bhad292.

Muñoz-Arnaiz, R., Makarova, J., Makarov, V.A., Herreras, O. (2025). Brain sources composing irregular field potentials have unique temporal signatures. Cereb Cortex, 35, bhaf135. doi:10.1093/cercor/bhaf135.

Nasretdinov, A., Vinokurova, D., Lemale, C.L., Burkhanova-Zakirova, G., Chernova, K., Makarova, J., Herreras, O., Dreier, J.P., Khazipov, R. (2023). Diversity of cortical activity changes beyond depression during spreading depolarizations. Nat Commun, 14, 7729. 10.1038/s41467-023-43509-3.

Ness, T.V., Remme, M.W., Einevoll, G.T. (2016). Active subthreshold dendritic conductances shape the local field potential. J Physiol, 594, 3809–3825. doi: 10.1113/JP272022.

Ness, T.V., Tetzlaff, T., Einevoll, G.T., Dahmen, D. (2025). On the validity of electric brain signal predictions based on population firing rates. PLoS Comput Biol, 21(4), e1012303. 10.1371/journal.pcbi.1012303.

Nunez, P.L., Srinivasan, R. (2006) Electric Fields in the Brain. The Neurophysics of EEG OUP.

Pincus, S.M. (1991). Approximate entropy as a measure of system complexity. Proc Nat Acad Sci, 88, 2297–2301. 10.1073/pnas.88.6.2297.

Reimann, M.W., Anastassiou, C.A., Perin, R., Hill, S.L., Markram, H., Koch, C. (2013). A biophysically detailed model of neocortical local field potentials predicts the critical role of active membrane currents. Neuron, 79, 375–390. doi: 10.1016/j.neuron.2013.05.023.

Rall, W., Shepherd, G.M. (1968). Theoretical reconstruction of field potentials and dendrodendritic synaptic interactions in olfactory bulb. J Neurophysiol, 31, 884–915. doi: 10.1152/jn.1968.31.6.884

Richman, J.S., Moorman, J.R. (2000). Physiological time-series analysis using approximate entropy and sample entropy. Am J Physiol-Heart Circ Physiol, 278, H2039–H2049. 10.1152/ajpheart.2000.278.6.H2039

Robinson, P.A., Rennie, C.J., Rowe, D.L., O’Connor, S.C., Wright, J.J., Gordon, E., Whitehouse, R.W. (2003). Neurophysical modeling of brain dynamics. Neuropsychopharmacology, 28(S1), S74–9. doi: 10.1038/sj.npp.1300143.

Rosso, O.A., Martin, M.T., Figliola, A., Keller, K., Plastino, A. (2006). EEG analysis using wavelet-based information tools. J Neurosci Meth, 153, 163–182. 10.1016/j.jneumeth.2005.10.009

Sanz Leon, P., Knock, S.A., Woodman, M.M., Domide, L., Mersmann, J., McIntosh, A.R., Jirsa, V. (2013). The Virtual Brain: a simulator of primate brain network dynamics. Front Neuroinform, 7, 10. doi: 10.3389/fninf.2013.00010.

Schomburg, E.W., Fernández-Ruiz, A., Mizuseki, K., Berényi, A., Anastassiou, C.A., Koch, C., Buzsáki, G. (2014). Theta phase segregation of input-specific gamma patterns in entorhinal-hippocampal networks. Neuron, 84, 470–485. doi: 10.1016/j.neuron.2014.08.051

Stam, C.J. (2005). Nonlinear dynamical analysis of EEG and MEG: review of an emerging field. Clin Neurophysiol, 116, 2266–2301. doi: 10.1016/j.clinph.2005.06.011

Telenczuk, M., Telenczuk, B., Destexhe, A. (2020a). Modelling unitary fields and the single-neuron contribution to local field potentials in the hippocampus. J Physiol, 598, 3957–3972. 10.1113/JP279452

Telenczuk, B., Telenczuk, M., Destexhe, A. (2020b). A kernel-based method to calculate local field potentials from networks of spiking neurons. J Neurosci Meth, 344, 108871. 10.1016/j.jneumeth.2020.108871

Tenke, C.E., Kayser, J. (2012). Generator localization by current source density (CSD): implications of volume conduction and field closure at intracranial and scalp resolutions. Clin Neurophysiol, 123, 2328–2345. doi: 10.1016/j.clinph.2012.06.005.

Tesler, F., Tort-Colet, N., Depannemaecker, D., Carlu, M., Destexhe, A. (2022). Mean-field based framework for forward modeling of LFP and MEG signals. Front Comput Neurosci, 16, 968278. doi: 10.3389/fncom.2022.968278

Tesler, F., Lorenzi, R.M., Ponzi, A., Casellato, C., Palesi, F., Gandolfi, D., Gandini Wheeler Kingshott, C.A.M., Mapelli, J., D’Angelo, E., Migliore, M., Destexhe, A. (2024). Multiscale modeling of neuronal dynamics in hippocampus CA1. Front Comput Neurosci, 18, 1432593. doi: 10.3389/fncom.2024.1432593.

Thivierge, J.P., Cisek, P. (2008). Nonperiodic synchronization in hetero-geneous networks of spiking neurons. J Neurosci, 28, 7968–7978. 10.1523/JNEUROSCI.0870-08.2008

Toharia, P., Robles, O.D., Fernaud-Espinosa, I., Makarova, J., Galindo, S.E., Rodriguez, A., Pastor, L., Herreras, O., DeFelipe, J., Benavides-Piccione, R. (2016). Pyramidal Explorer: A new interactive tool to explore morpho-functional relations of pyramidal neurons. Front Neuroanat, 9,159. doi: 10.3389/fnana.2015.00159

Torres, D., Makarova, J., Ortuño, T., Benito, N., Makarov, V.A., Herreras, O. (2019). Local and volume-conducted contributions to cortical field potentials. Cereb Cortex, 29, 5234–5254. doi: 10.1093/cercor/bhz061

Varona, P., Ibarz, J.M., López-Aguado, L., Herreras, O. (2000). Macroscopic and subcellular factors shaping CA1 population spikes. J Neurophysiol, 83, 2192–2208. 10.1152/jn.2000.83.4.2192

Yuste, R., Cossart, R., Yaksi, E. (2024). Neuronal ensembles: Building blocks of neural circuits. Neuron, 112, 875–892 doi: 10.1016/j.neuron.2023.12.008

## Supplementary References

Delgado-Bonal A, Marshak A. Approximate entropy and sample entropy: A comprehensive tutorial. Entropy 21:541 (2019). 10.3390/e21060541

López-Madrona VJ, et al. Different theta frameworks coexist in the rat hippocampus and are coordinated during memory-guided and novelty tasks. eLife 9, e57313 (2020). doi: 10.7554/eLife.57313.

Makarov VA, Muñoz R, Herreras O, and Makarova J. Correlation dimension of high-dimensional and high-definition experimental time series. Chaos 33, 123114 (2023). doi: 10.1063/5.0168400.

Montgomery DC, Runger GC. (2014). Applied Statistics and Probability for Engineers (6th ed.). John Wiley & Sons.

Mormann F et al. On the predictability of epileptic seizures. Clin Neurophysiol. 116:569–587 (2005). 10.1016/j.clinph.2004.08.025.

Rényi A. (1961). On measures of entropy and information. In: Contributions to the theory of statistics. Vol 4. Berkeley, CA: University of California Press, 547–562.

Sauer T, Yorke JA, Casdagli M. Embedology J Stat Phys. 65:579–616 (1991). 10.1007/BF01053745.

Takens, F. 1981. Detecting strange attractors in turbulence. In: Rand DA, Young L-S, editors. Dynamical Systems and Turbulence. Lecture Notes in Mathematics, vol 898. Springer, Berlin, Heidelberg. 10.1007/BFb0091924

